# Loss of nuclear-cytoplasmic compartmentalization is a hallmark of aging in podocytes

**DOI:** 10.64898/2026.04.29.721580

**Authors:** Mohamed Hamed, Yingying Gao, Frank Stein, Grishal Vijay, Thilo Stausberg, Ramona Jühlen, Ina V. Martin, Ute Raffetseder, Eleni Stamellou, Rafael Kramann, Tammo Ostendorf, Wolfram Antonin

## Abstract

The nuclear envelope, with its nuclear pore complexes, establishes a selective barrier that maintains the spatial organization of the cellular proteome. Whether this barrier remains intact in long-lived postmitotic cells during aging is largely unknown. Here, we show that aging podocytes experience a progressive loss of nuclear-cytoplasmic compartmentalization. Using subcellular proteomics and quantitative imaging, we identify a global redistribution of proteins between the nucleus and cytoplasm, affecting key regulators of RNA processing, chromatin organization, and cellular metabolism. Loss of compartmentalization is accompanied by mitochondrial dysfunction, increased reactive oxygen species production, and aberrant nuclear accumulation of YAP1, linking nuclear barrier failure to metabolic and transcriptional dysregulation. These findings identify nuclear envelope dysfunction as a driver of proteome disorganization and cellular decline in aging podocytes and suggest that loss of nuclear-cytoplasmic compartmentalization represents a general mechanism contributing to dysfunction in long-lived cells.

## Introduction

The establishment of subcellular compartmentalization is regarded as a pivotal event in the evolution of organisms, marking the transition from the simpler structures of prokaryotic cells to the more complex eukaryotic cells (1). This transformation involved the acquisition of several key features, including a nucleus, an endomembrane system, a cytoskeleton, and energy-producing organelles, through endosymbiosis. Intricate compartmentalization within the cell improves the efficiency of cellular reactions and processes by creating suitable microenvironments for biochemical activities. By separating incompatible reactions into distinct microenvironments, this organization enables multiple biochemical pathways to occur simultaneously (2). Additionally, it enhances metabolic efficiency by gathering substrates, enzymes, and cofactors in specific areas, reducing the loss of intermediates. Furthermore, it enables precise spatial control over signaling by guiding specific pathways to different organelles or subdomains (3). In eukaryotes, this compartmentalization allows for a level of gene expression regulation that is not present in prokaryotes, as it spatially and temporally separates transcription from translation (4).

Because of its vital importance, the loss of subcellular compartmentalization is directly associated with various disease types, as it interrupts the spatial regulation of signaling, gene control, and proteostasis (5–7). Similarly, this loss may arise from the decline in cellular function observed during aging, which is often linked to the accumulation of damage. This phenomenon could be especially important in long-lived postmitotic cells, such as neurons (8) or podocytes, which create the filtration barrier in the kidney. Accordingly, podocyte injury, characterized by loss of cell function and eventual cell depletion, has been associated with the decline in kidney function during aging (9–13). It is therefore possible, but not yet investigated, that a loss of subcellular compartmentalization may also contribute to podocyte injury and functional loss.

In eukaryotic cells, the cytoplasm and nucleus represent the two major compartments, separated by the nuclear envelope, which functions as a selective diffusion barrier. Embedded in this envelope are nuclear pore complexes (NPCs) that facilitate efficient communication between the cytoplasm and the nucleoplasm (14). NPCs carry out the selective, regulated exchange of proteins and ribonucleoprotein particles: nuclear transport receptors (NTRs) recognize nuclear localization or export signals on cargo proteins, form transport-competent complexes, and mediate their translocation through NPCs (15). The small GTP-binding protein RAN provides directionality to nucleo-cytoplasmic transport: on the nuclear side, RAN-GTP prompts the disassembly of import complexes and enables export receptors to bind their cargo and form export complexes in the nucleus, which can then pass through NPCs (15, 16). At the same time, NPCs establish a selective permeability barrier that restricts the passive diffusion of most macromolecules. This barrier is mainly formed by intrinsically disordered FG repeat nucleoporins lining the central channel, functioning as a size and hydrophobicity dependent filter. Consequently, proteins smaller than approximately 30–40 kDa diffuse relatively freely through the pore, whereas larger macromolecules are excluded unless actively transported by nuclear transport factors (15, 16).

Here, we investigate how aging affects nuclear-to-cytoplasmic compartmentalization in podocytes, specialized long-lived postmitotic cells that are essential for maintaining the glomerular filtration barrier and normal kidney function. Combining subcellular fractionation, quantitative proteomics, and imaging approaches, we show that aging podocytes exhibit a marked redistribution of proteins between the nucleus and the cytoplasm. This shift is accompanied by mitochondrial dysfunction, increased production of reactive oxygen species, and enhanced nuclear accumulation of YAP1, consistent with activation of mechanical stress signaling.

## Results

### Aging mice exhibit progressive podocyte loss and early focal segmental glomerulosclerosis

To investigate the impact of aging on podocytes and kidney function, we established a murine aging model. Mice were stratified into three age groups: 11 weeks (young), 35 weeks (middle-aged), and 80 weeks (aged), corresponding approximately to 21, 41, and 90 years in humans, respectively (17).

Previous studies have established a strong association between aging, podocyte loss, and the development of focal segmental glomerulosclerosis (FSGS) (12, 13, 18). We therefore characterized our aging mouse model by quantifying podocyte numbers using 3D two-photon microscopy (Fig. 1a) following doxycycline-induced eGFP labeling of podocyte nuclei. This analysis revealed an age-dependent decline in podocyte numbers. Specifically, podocyte counts dropped to a mean of 48 per glomerulus at 80 weeks, compared with 71 at 35 weeks and 73 at 11 weeks. These findings were consistent across glomeruli within each age group and indicate a progressive, age-associated loss of podocytes.

**Figure 1:**
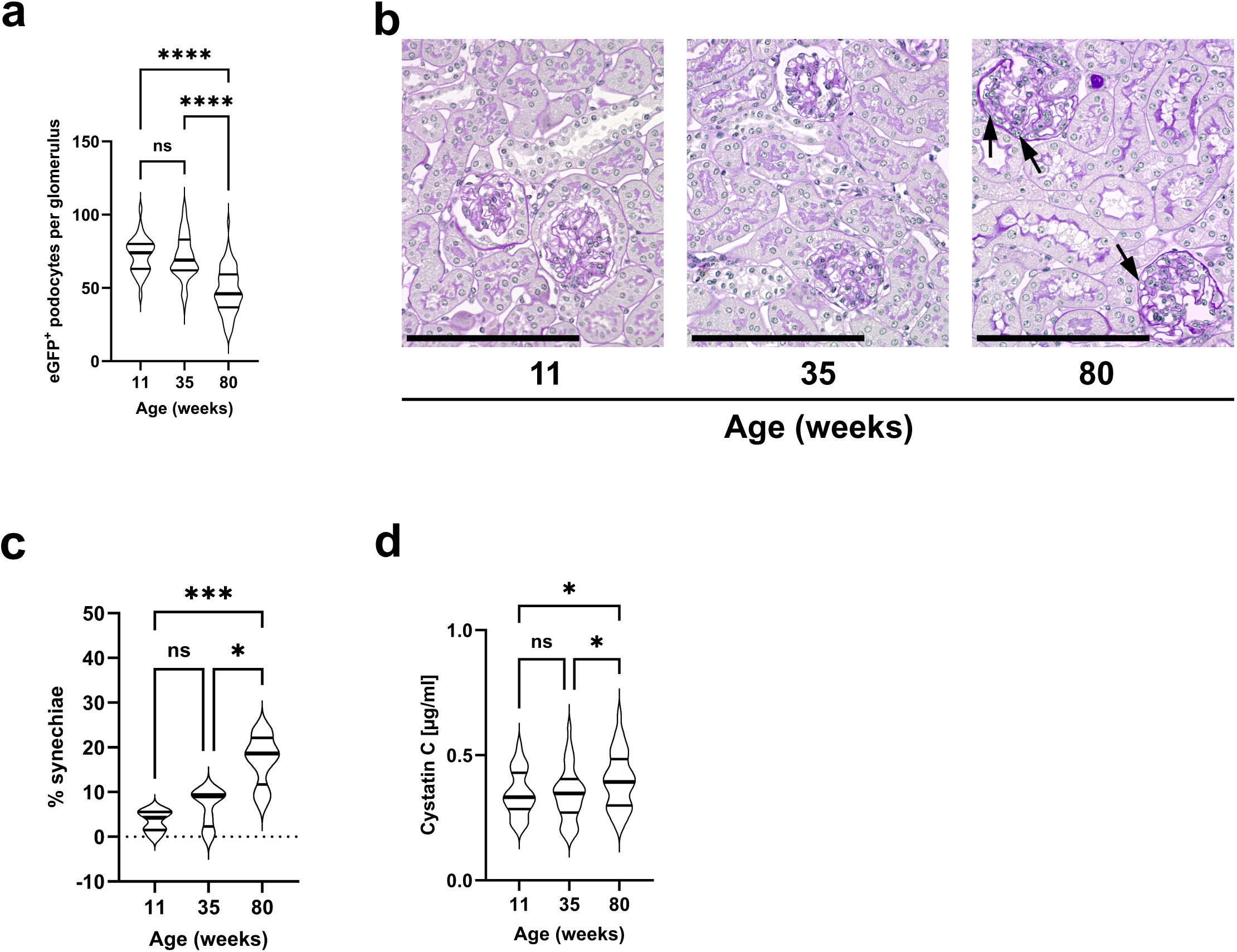
Aging drives podocyte loss, initial glomerular damage, and decline in kidney function in mice. (a) Quantification of eGFP^+^ podocytes using a custom pipeline to measure the number of podocytes expressing a GFP signal in the nucleus. 11-week group (n = 34 podocytes), 35-week group (n = 75 podocytes), 80-week group (n = 62 podocytes). Data is shown as violin blots; thick lines are the medians, thin lines the 25% and 75% quartiles. ns: non-significant, ****P< 0.0001. Ordinary one-way ANOVA, Fisher’s LSD test, multiple comparisons; 11 weeks vs. 35 weeks, p=0.5880, 11 weeks vs. 80 weeks, ****P<0.0001, 35 weeks vs. 80 weeks, ****P<0.0001. (b) Representative PAS-stained kidney sections from mice of different ages illustrating age-associated glomerular changes. Kidneys from 80-week-old mice display glomeruli with adhesions of the glomerular tuft to Bowman’s capsule, forming synechiae (right panel, black arrows), consistent with an early stage of focal segmental glomerulosclerosis (FSGS) and indicative of podocyte injury and activation of parietal epithelial cells. Scale bar: 100 µm. (c) Quantification of glomerular synechiae in the three age groups: 11 weeks (n = 7 mice), 35 weeks (n = 8 mice), and 80 weeks (n = 8 mice). Violin blots; thick lines are the medians, thin lines the 25% and 75% quartiles. ns: non-significant, *P < 0.05, ***P < 0.001. Kruskal-Wallis non-parametric test, one-way ANOVA, uncorrected Dunn’s test, multiple comparisons; 11 weeks vs. 35 weeks, P=0.2134, 11 weeks vs. 80 weeks, ***P=0.0003, 35 weeks vs. 80 weeks, *P=0.0142. (d) Serum Cystatin C levels at 11, 35, and 80 weeks (n = 34 mice per condition). Violin blots; thick lines are the medians, thin lines the 25% and 75% quartiles. ns: non-significant, *P < 0.05. Ordinary one-way ANOVA, Fisher’s LSD test, multiple comparisons; 11 weeks vs. 35 weeks, P=0.8657, 11 weeks vs. 80 weeks, *P=0.0396, 35 weeks vs. 80 weeks, *P=0.0264.

Along with aging, we observed a marked increase in glomerular synechiae (Fig. 1b-c), adhesions of the glomerular tuft to Bowman’s capsule, and an early hallmark of FSGS. By 80 weeks, the incidence of synechiae had risen to 17.2%, substantially higher than at 35 weeks (6.7%) and 11 weeks (3.6%).

Consistent with progressive podocyte loss and early FSGS lesions in aged mice, kidney function declined with age, as indicated by increased serum Cystatin C levels, particularly in the 80-week group (Fig. 1d). These findings collectively suggest a strong link between aging, podocyte depletion, and FSGS development (12, 13, 18), confirming that our aging model provides an appropriate system to investigate podocyte pathophysiology, with a particular focus on cellular compartmentalization and NPC function during aging.

### Nuclear pore complex barrier function deteriorates in aging podocytes

Because aging has been associated with a decline in NPC function, at least in neurons (19), and mutations in nucleoporins can result in FSGS (20, 21), we tested whether NPC function is specifically impaired in podocytes from aged mice. Podocytes were isolated from mice at 11, 35, and 80 weeks. Glomeruli were obtained from kidneys and enzymatically digested to yield individual podocytes, which were purified by FACS based on nuclear eGFP expression driven by a podocyte-specific promoter (see Methods for details). The plasma membranes of the isolated podocytes were permeabilized, and the cells were incubated with dextrans of different sizes (10, 40, and 70 kDa). NPCs act as size-dependent diffusion barriers for macromolecules between the cytoplasm and nucleoplasm (16). Smaller molecules can pass freely, whereas larger ones are excluded unless actively transported. In podocytes from young mice, fluorescently labeled dextrans of 10 kDa readily penetrate the nucleus, whereas 40 kDa and 70 kDa dextrans are excluded (Fig. 2a-c, 11 weeks panel, d). By contrast, nuclei from podocytes of 80-week-old mice are labeled by both 40 kDa and 70 kDa dextrans, indicating a decline in NPC barrier function (Fig. 2a-c, 80 weeks panel, d). Nuclei from 35-week-old mice show increased leakage of 40 kDa and 70 kDa dextrans compared with 11-week-old mice, but less than nuclei from 80-week-old mice (Fig. 2a-c, 35 weeks panel, d). While we cannot completely rule out effects of podocyte isolation on NPC integrity, these results suggest that NPCs in older podocytes are more prone to leakiness than those in younger cells, which maintain greater resilience.

**Figure 2:**
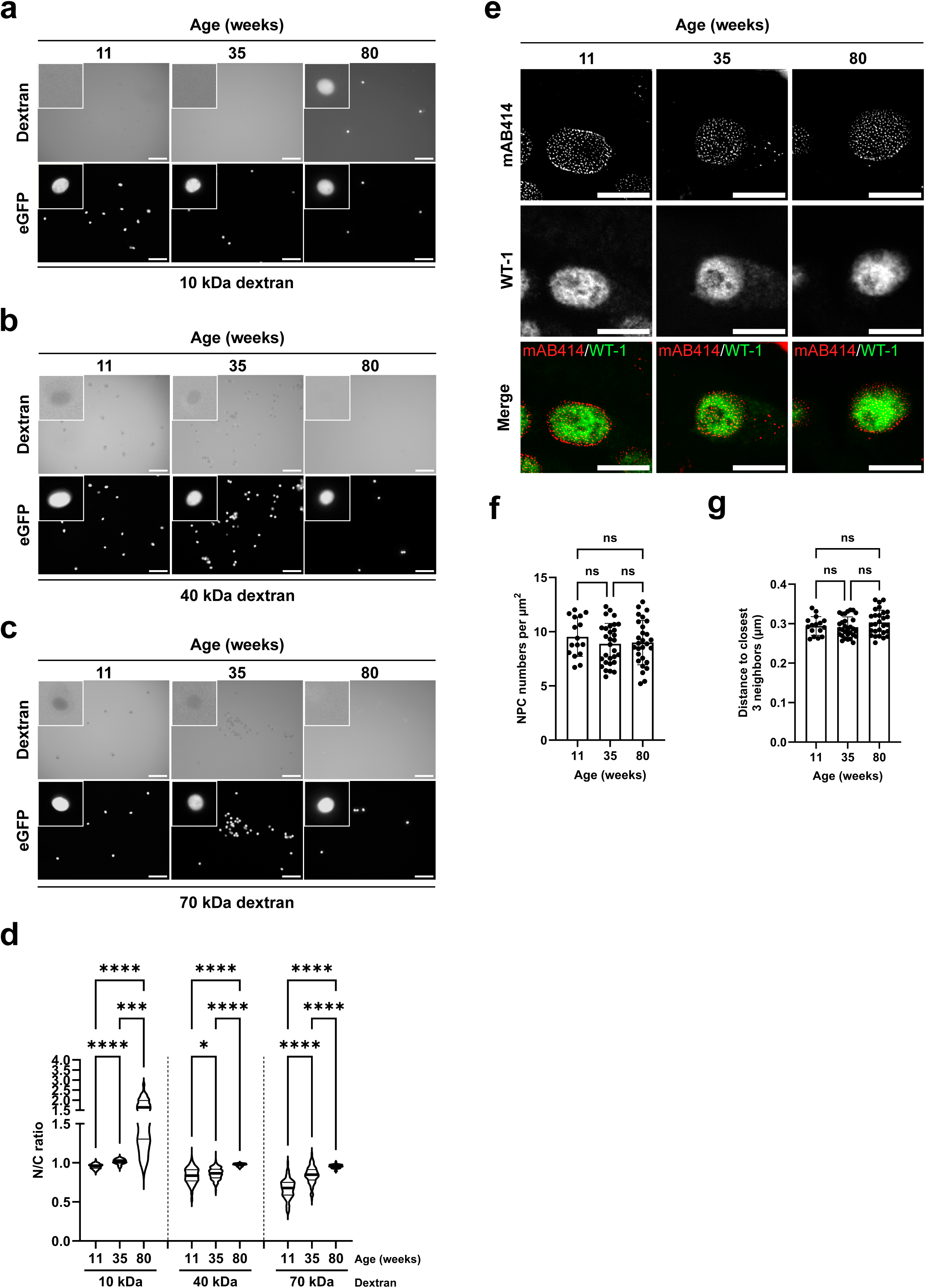
Aged podocytes exhibit altered nuclear permeability but maintain constant NPC density. (**a-c**) Representative fluorescence images of eGFP primary podocytes from 11-, 35-, and 80-week-old mice, permeabilized with 0.2% NP-40 and incubated with Texas Red-labeled 10 kDa (**a**), 40 kDa (**b**), or 70 kDa (**c**) dextran. Podocyte nuclei are identified by eGFP (lower panels). Scale bar, 100 μm. Insets show high-magnification views of individual nuclei. (d) Quantification of dextran permeability in **a-c**, expressed as the nuclear-to-cytoplasmic fluorescence intensity ratio (n = 3 biological replicates for 11 weeks, from 6 mice; n = 2 for 35 weeks, from 5 mice and n=2 for 80 weeks, from 6 mice). Thick lines indicate medians; thin lines show the 25% and 75% quartiles. *P < 0.05, ***P < 0.001, ****P < 0.0001. Kruskal-Wallis non-parametric test, one-way ANOVA, uncorrected Dunn’s test, multiple comparisons; 10 kDa, 11 weeks vs. 35 weeks, ****P<0.0001, 11 weeks vs. 80 weeks, ****P<0.0001, 35 weeks vs. 80 weeks, ***P=0.0006; 40 kDa, 11 weeks vs. 35 weeks, *P=0.0269, 11 weeks vs. 80 weeks, ****P<0.0001, 35 weeks vs. 80 weeks, ****P<0.0001; 70 kDa, ****P<0.0001. (e) Representative STED images of podocytes, identified by the podocyte-specific nuclear WT-1 staining (green overlay), displaying individual NPCs labeled with the antibody mAB414 (red overlay). Scale bar, 5 μm. (**f, g**) Quantification of NPC density (**f**) and the distance to the three nearest neighboring NPCs (**g**) from samples imaged as in E. n = 16 podocytes for 11 weeks, from 4 mice; n = 30 podocytes for 35 weeks, from 6 mice, and n=28 podocytes for 80 weeks, from 4 mice. Data are presented as means ± SD. ns: non-significant. Ordinary one-way ANOVA, Fisher’s LSD test, multiple comparisons were used for (**f**), Kruskal-Wallis non-parametric test, one-way ANOVA, uncorrected Dunn’s test, and multiple comparisons were used for (**g**).

We next examined whether the decline in NPC function is accompanied by a loss of NPCs in aging mouse podocytes. Kidney tissues were stained with mAB414, an antibody that detects multiple nucleoporins within the NPC and is widely regarded as a reliable NPC marker (22), alongside nuclear WT-1 to identify podocytes (Fig. 2e). High-resolution stimulated emission depletion (STED) microscopy showed no significant changes in either the NPC density (Fig. 2f) or the average distance of each NPC to its three closest neighbors (Fig. 2g) across the three age groups. These results indicate that both NPC numbers per nucleus and NPC density remain largely constant in podocytes during aging.

### Subcellular proteomics reveals changes in nuclear□to□cytoplasmic ratios in aging podocytes

Because NPCs are crucial for establishing and maintaining nuclear cytoplasmic compartmentalization, we investigated whether an age related decline in NPC mediated exclusion affects the localization of nuclear and cytoplasmic proteins in mice. Podocytes were isolated from three age groups (11, 35, and 80 weeks), and cytoplasmic and nucleoplasmic fractions were prepared from each sample (Fig. 3a, Supplemental Fig. 1a) for quantitative mass spectrometry analysis (see Methods). Across all biological replicates within each age group, a total of 3,583 proteins were reliably identified. For each protein, we calculated the nucleoplasmic to cytoplasmic (N/C) ratio: proteins with a ratio > 1.0 were classified as nuclear, whereas those with a ratio < 1.0 were classified as cytoplasmic. Log fold changes of these ratios were visualized in a correlation map (Fig. 3b) to illustrate how subcellular distribution changes or remains stable during aging. Proteins predominantly nuclear are shown in red, and predominantly cytoplasmic proteins are shown in blue. Based on their localization at the three time points (11, 35, and 80 weeks), proteins were grouped into four clusters. Cluster I includes 554 proteins that are mainly cytoplasmic at 11 weeks and shift to mainly nuclear by 80 weeks (485 are already predominantly nuclear at 35 weeks). Cluster II includes 264 proteins that are mainly nuclear at 11 weeks and become mostly cytoplasmic by 80 weeks (228 are already predominantly cytoplasmic at 35 weeks). Cluster III comprises 1289 proteins that remain nuclear at all-time points, and Cluster IV comprises 1476 proteins that remain cytoplasmic at all-time points.

**Figure 3:**
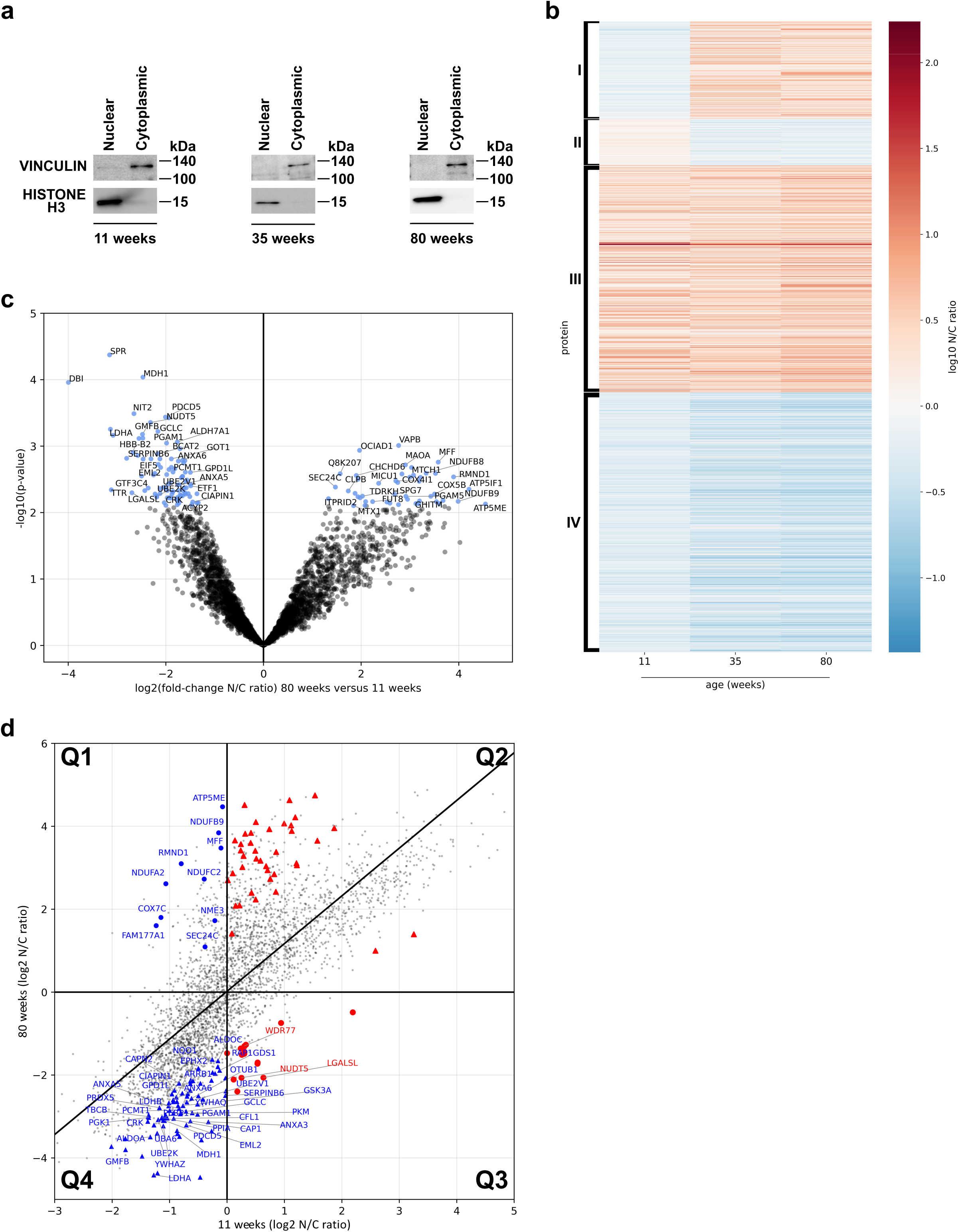
Aging drives protein redistribution between the nucleus and cytoplasm in podocytes. (a) Western blots of nuclear and cytoplasmic fractions from primary podocytes isolated from 11-, 35-, and 80-week-old mice, using VINCULIN as a cytoplasmic marker and HISTONE H3 as a nuclear marker. (b) Heat map showing age-related changes in subcellular protein distribution in podocytes from 11, 35, and 80-week-old mice, displaying the log10-transformed nuclear-to-cytoplasmic ratio for each protein. Proteins are grouped into four clusters based on their localization patterns: cluster I, cytoplasmic to nuclear shift; cluster II, nuclear to cytoplasmic shift; cluster III, stable nuclear; and cluster IV, stable cytoplasmic localization. Data are Z-score normalized across all samples; color intensity indicates relative protein abundance. Each column represents the average of biological replicates (n = 3 for 11 weeks, from 20 mice; n = 3 for 35 weeks, from 18 mice; n = 2 for 80 weeks, from 46 mice). (c) Volcano plot illustrating differential subcellular protein distribution in podocytes from 80-week-old versus 11-week-old mice. The x-axis depicts the log2-fold change in the nuclear-to-cytoplasmic ratio, while the y-axis shows the −log10 (p-value). Proteins not meeting significance thresholds (P > 0.05, absolute fold change < 1.5, false discovery rate (FDR) > 0.2) are shown in grey; significantly altered proteins (P < 0.05, absolute fold change > 1.5 and false discovery rate (FDR) < 0.2) are in blue. (d) Scatter plot comparing subcellular protein distribution in podocytes from 11 versus 80-week-old mice. The x-axis displays the log2-transformed nuclear-to-cytoplasmic ratio of podocytes from 11-week-old mice, and the y-axis displays the log2-transformed nuclear-to-cytoplasmic ratio from 80-week-old mice. The axes indicate equal nuclear and cytoplasmic distributions, with the diagonal line representing no change with age. Significantly altered proteins (P < 0.05, absolute fold change > 1.5 and false discovery rate (FDR) < 0.2) are labeled based on localization dynamics: nuclear to cytoplasmic (red circles), cytoplasmic to nuclear (blue circles) shifts, stable cytoplasmic (blue triangles), or stable nuclear (red triangles) localization. The quadrants are labeled Q1, Q2, Q3, and Q4.

To further illustrate how subcellular distribution changes in aging podocytes, we plotted the log_2_-fold change of the nuclear-to-cytoplasmic ratio between 80 and 11 weeks in a volcano plot (Fig. 3c). Proteins showing significant changes (n=132, p-value < 0.05) are highlighted in blue. Of these, 89 proteins shifted toward a more cytoplasmic localization, indicating a decrease in the N/C ratio (FC < 0.5, left side of the plot), whereas 43 proteins shifted toward a more nuclear localization, with increased N/C ratios (FC > 1.5, right side of the plot).

To visualize sub-cellular distribution over time, we plotted the N/C ratio of each protein at 11 weeks against its N/C ratio at 80 weeks as a fold-change correlation (Fig. 3d). Proteins in quadrant 1 are mainly cytoplasmic at 11 weeks and shift to predominantly nuclear localization by 80 weeks, corresponding to cluster I in Fig. 3b; 10 of these proteins reach significance and are labeled as blue circles. Conversely, quadrant 3 contains proteins that are mainly nuclear at 11 weeks and shift to predominantly cytoplasmic localization by 80 weeks, corresponding to cluster II in Fig. 3b, with 13 proteins reaching statistical significance (red circles). Quadrants 2 and 4 contain proteins that maintain predominantly nuclear or cytoplasmic localization over time.

A notable subset of nuclear proteins that change their subcellular distribution to the cytoplasm with age are involved in chromatin remodeling, including NUDT5 and WDR77 (Fig. 3d). Additional chromatin remodeling factors that show similar localization changes but do not reach the significance threshold in Fig. 3c/d include ACTL6A, CFDP1, MIER1, RBBP4, RUVBL1, and SMYD5. Furthermore, several mRNA-processing factors, including HNRNPH2, MBNL1, and ZCCHC8, also exhibit increased cytoplasmic localization in aged podocytes.

Similarly, the small GTPase RAN, which governs the directionality of nuclear transport (15, 16), shifts toward the cytosol in aged podocytes (Supplemental Fig. 2). Likewise, specific nuclear-import receptors, importin α1 and the importin β family members IPO4, IPO5, IPO7, and IPO9, which normally cycle between the cytoplasm, where they bind cargo, and the nucleus, show a significant reduction in nuclear localization. A similar pattern is observed for the export receptors XPO1, XPO5, and XPO7, which are usually nuclear, bind cargo with RAN-GTP, and shuttle between compartments. They exhibit an even lower nuclear presence in aged cells. Together, these findings suggest a decline in NPC functionality, consistent with the reduced dextran exclusion capacity observed in the functional assays (Fig. 2a-d).

A subset of cytoplasmic proteins that relocalize to the nucleus in aged podocytes includes key components of vesicular trafficking, such as the significantly altered proteins FAM177A1 and SEC24C (Fig 3d). Additional factors include GRIPAP1, RABL3, TM9SF3, VAMP4, and dynamin II (DNM2), which is involved in membrane remodeling and cell motility. Notably, a large group of mitochondrial proteins shows increased nuclear localization in aged podocytes, including subunits of the respiratory chain and ATP synthase (ATP5ME, COX7C, NDUFA2, NDUFB9, and NDUFC2), as well as mitochondrial fission and fusion factors (MFF and NME3), and RMND1, a protein critical for mitochondrial DNA synthesis (Fig 3d). Additional proteins below the significance threshold include mitochondrial ribosomal proteins (MRPL2, MRPL27, MRPL32, MRPL41, MRPL54, and MRPS16), as well as auxiliary factors like ECSIT, the antioxidant enzyme GPX4, and further components of the respiratory chain (ATP5PD, ATP5PF, COX6B1, NDUFC1, NDUFS6, NDUFV2, NDUFV3, and UQCRH). Together, the cytoplasmic relocation of chromatin remodeling and RNA processing factors, along with the nuclear enrichment of vesicular trafficking regulators and mitochondrial components, highlights a broad disruption of intracellular compartmentalization in aged podocytes. These changes are consistent with impaired NPC selectivity and increased permeability, as supported by dextran-exclusion assays.

Expectably, many proteins change their overall abundance with aging without a corresponding shift in their nuclear-cytoplasmic distribution, indicating stable relative localization between compartments. These include cytoplasmic proteins (Fig. 3d, quadrant 4) involved in general cellular signaling (ARRB1, CRK, GSK3A, PEBP1, RAP1GDS1, YWHAQ, and YWHAZ), cytoskeletal organization (ANXA3, ANXA5, ANXA6, CAP1, CAPN2, CFL1, EML2, GMFB, PPIA, and TBCB), stress response (EPHX2, GCLC, NQO1, and PRDX5), energy metabolism (ALDOA, ALDOC, GPD1L, LDHA, LDHB, PGAM1, PGK1, AND PKM), apoptosis (CIAPIN1 and PDCD5) and protein folding (UBA6, UBE2K, UBE2V1, OTUB1, and PCMT1). Several of these proteins have established roles in podocyte biology. For example, loss of PRDX5 increases reactive oxygen species (ROS) production, whereas NQO1 deficiency exacerbates podocyte injury (23, 24). Similarly, impairment of 14-3-3 proteins (YWHAQ and YWHAZ) and annexins (ANXA3, ANXA5, and ANXA6) reduces the stability of plasma membrane-actin interactions and cytoskeletal integrity, thereby increasing susceptibility to mechanical stress (25–27). In addition, CFL1 is essential for maintaining actin dynamics and cytoskeletal organization in podocytes (28).

Similarly, a subset of proteins shows increased nuclear abundance while remaining predominantly confined to the nuclear compartment. This group includes chromatin regulators. ATP-dependent chromatin-remodeling components include AZ1B, BAZ2A, BPTF, CHD2, RUVBL2, SMARCA5, SMARCD1, SRCAP, and VPS72. In addition, numerous epigenetic modifiers and chromatin-associated proteins are present, such as BANF1, BRD2, BRD3, BRD4, CBX4, CBX8, CTCF, DEK, EHMT1, EHMT2, GATAD2A, H2AZ1, HCF1, HMGB1, HMGB3, HMGN1, HMGN2, KAT8, KDM1A, MACROH2A1, MACROH2A2, MBD1, MBD2, MECP2, and SMCHD1. These factors regulate histone modifications, higher-order chromatin organization, and transcription, indicating a pronounced remodeling of chromatin-regulatory programs in aged podocytes.

### Nucleo-cytoplasmic ratios of candidate proteins change during aging in podocytes

To validate the age-dependent shifts in nucleocytoplasmic distribution observed in the proteomic screen, we employed an independent quantitative imaging approach (29). Kidney sections were stained with validated antibodies for immunofluorescence analysis and co-stained with WT-1 and NESTIN as markers of podocyte nuclei and cytoplasm, respectively. For each podocyte, fluorescence intensities of the candidate proteins were measured in the nucleus and cytoplasm, allowing calculation of the nucleocytoplasmic (N/C) ratio. Based on the proteomic data, we selected two proteins that translocate from the nucleus to the cytoplasm with age, SERPINB6 and LGALSL (Fig. 4), and four proteins that translocate from the cytoplasm to the nucleus with age, SEC24C, NDUFA2, NDUFS1, and ATP5PD (Fig. 5 and Supplemental Fig. 3).

**Figure 4:**
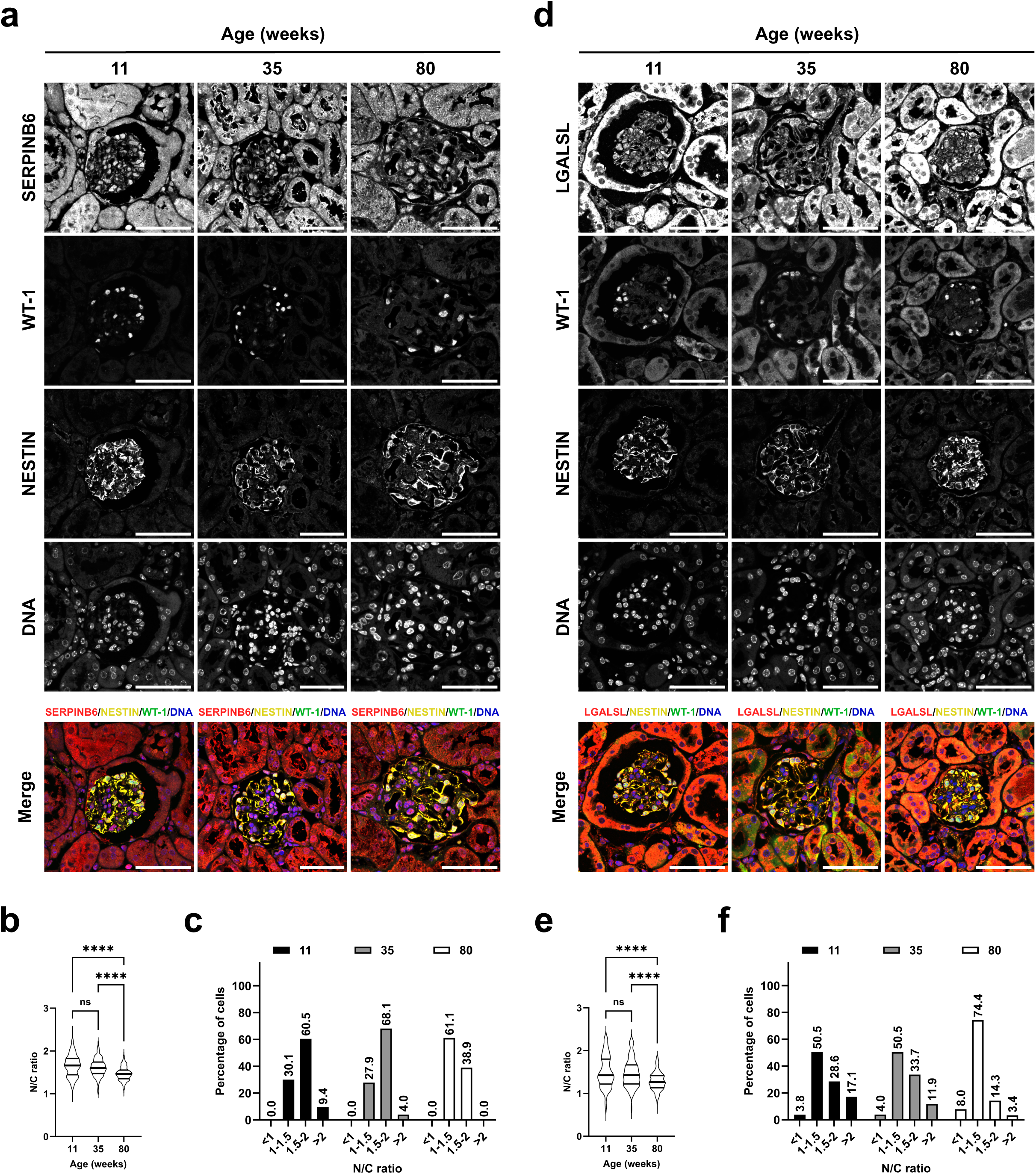
Age-dependent SERPINB6 and LGALSL localization in podocytes. (a) Representative confocal images of kidney sections from 11, 35, and 80-week-old mice stained for SERPINB6 (red), with NESTIN marking the podocyte cytoplasm (yellow), WT-1 marking the podocyte nucleus (green), and DAPI labeling DNA (blue). Scale bar, 50 µm. (b) Quantification of the nuclear-to-cytoplasmic ratio of SERPINB6 staining intensity as shown in (**a**). Thick lines represent medians, thin lines indicate the 25% and 75% quartiles, with n = 233 podocytes analyzed for 11 weeks from 6 mice, n = 323 for 35 weeks from 6 mice, and n = 352 for 80 weeks from 7 mice. ns: non-significant, ****P < 0.0001. Kruskal-Wallis non-parametric test, one-way ANOVA, uncorrected Dunn’s test, multiple comparisons; 11 weeks vs. 35 weeks, P=0.2372, 11 weeks vs. 80 weeks, ****P < 0.0001, 35 weeks vs. 80 weeks, ****P < 0.0001. (c) Distribution of podocytes into four nuclear-to-cytoplasmic ratio categories (NCs < 1, 1–1.5, 1.5–2, > 2), with percentages displayed above each column. (d) Representative confocal images of LGALSL staining (red) in kidney sections from 11, 35, and 80-week-old mice, stained and co-stained as in (**a**). Scale bar, 50 µm. (e) Quantification of the nuclear-to-cytoplasmic ratio of LGALSL as shown in (**b**), with n = 98 podocytes for 11 weeks from 4 mice, n = 96 for 35 weeks from 6 mice, and n = 228 for 80 weeks from 6 mice. ns: non-significant, ****P< 0.0001. Ordinary one-way ANOVA, Fisher’s LSD test, multiple comparisons; 11 weeks vs. 35 weeks, p=0.4383, 11 weeks vs. 80 weeks, ****P<0.0001, 35 weeks vs. 80 weeks, ****P<0.0001. (f) Distribution of podocytes into four nuclear-to-cytoplasmic ratio categories as in (**c**).

**Figure 5:**
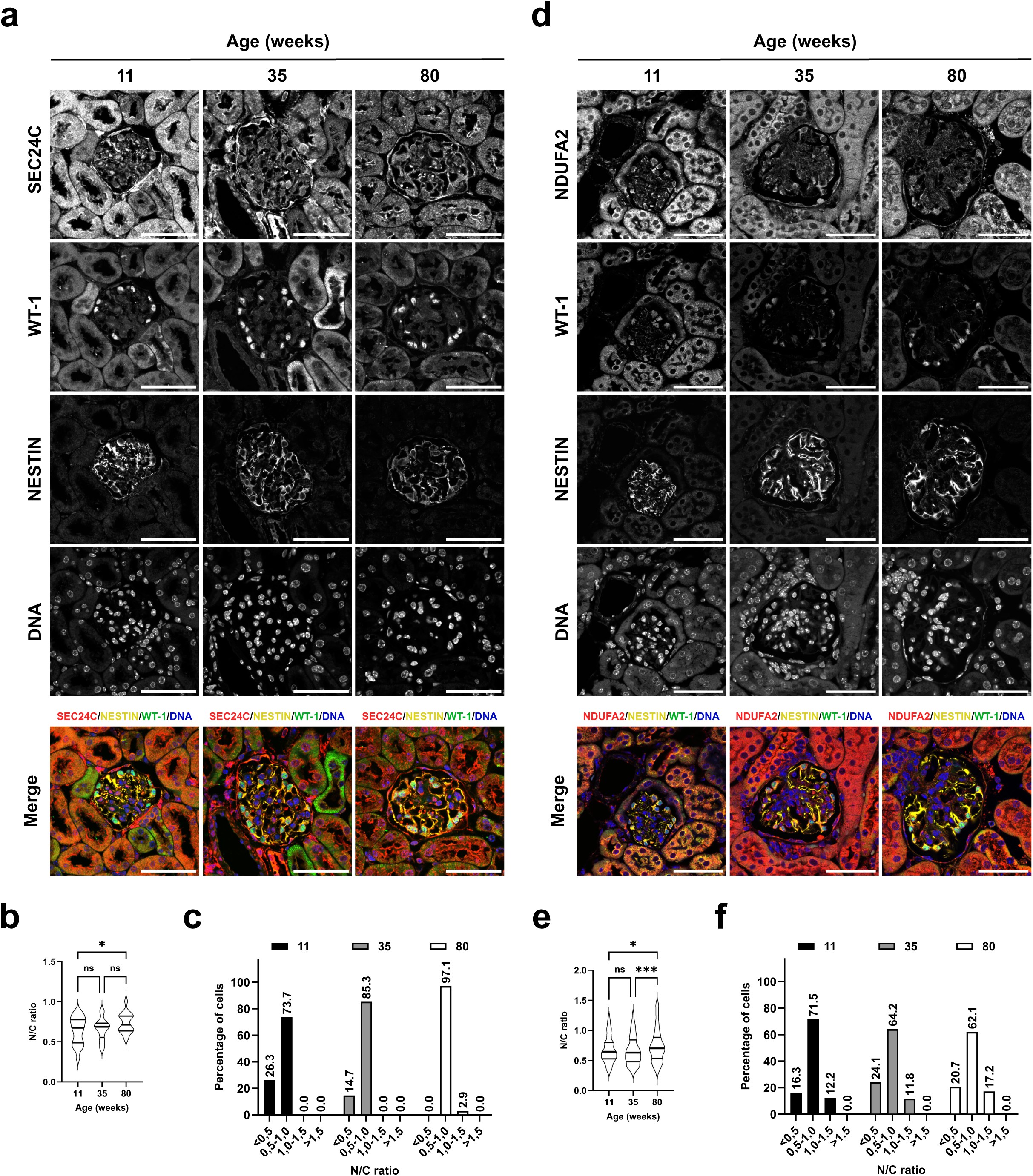
Age-dependent SEC24C and NDUFA2 localization in podocytes. (a) Representative confocal images of kidney sections from 11, 35, and 80-week-old mice stained for SEC24C (red), with NESTIN marking the podocyte cytoplasm (yellow), WT-1 marking the podocyte nucleus (green), and DAPI labeling DNA (blue). Scale bar, 50 µm. (b) Quantification of the nuclear-to-cytoplasmic ratio of SEC24C staining intensity as in (**a**). Thick lines indicate medians, thin lines show the 25% and 75% quartiles; n = 38 for 11 weeks from 6 mice, n = 34 for 35 weeks from 7 mice, and n = 34 for 80 weeks from 7 mice. ns: non-significant, *P < 0.05. Ordinary one-way ANOVA, Fisher’s LSD test, multiple comparisons; 11 weeks vs. 35 weeks, p=0.9356, 11 weeks vs. 80 weeks, *P=0.0172, 35 weeks vs. 80 weeks, P=0.0811. (c) Distribution of podocytes into four nuclear-to-cytoplasmic ratio categories (NCs: <0.5, 0.5–1.0, 1.0–1.5, >1.5), with percentages displayed above the corresponding columns. (d) Representative confocal images of NDUFA2 staining (red) in kidney sections from 11, 35, and 80-week-old mice, co-stained as in (**a**). Scale bar, 50 µm. (e) Quantification of the nuclear-to-cytoplasmic ratio of NDUFA2 as in (**b**); n = 195 for 11 weeks from 6 mice, n = 235 for 35 weeks from 4 mice, and n = 404 for 80 weeks from 7 mice. ns: non-significant, *P < 0.05, ***P < 0.001. Kruskal-Wallis non-parametric test, one-way ANOVA, uncorrected Dunn’s test, multiple comparisons; 11 weeks vs. 35 weeks, P=0.3605, 11 weeks vs. 80 weeks, *P=0.0199, 35 weeks vs. 80 weeks, ***P=0.0004. (f) Distribution of podocytes into four nuclear-to-cytoplasmic ratio categories as in (**c**).

SERPINB6, an intracellular serine protease inhibitor that localizes to both the nucleus and cytoplasm (30), showed an N/C ratio of 1.471 ± 0.160 at 80 weeks, compared to 1.614 ± 0.203 at 35 weeks and 1.655 ± 0.256 at 11 weeks (Fig. 4a, b). To further assess these changes, we stratified N/C ratios into four mutually exclusive categories: >2, indicating strong nuclear enrichment; 1.5–2.0, moderate nuclear enrichment; 1.0–1.5, approximately equal nuclear and cytoplasmic distribution; and <1, predominantly cytoplasmic localization. This classification showed a complete absence of cells with N/C ratios >2 at 80 weeks (0.0%), compared with 4.0% at 35 weeks and 9.4% at 11 weeks (Fig. 4c). Conversely, the proportion of cells with N/C ratios between 1.0 and 1.5 increased significantly, reaching 61.1% at 80 weeks compared with 27.9% at 35 weeks and 30.1% at 11 weeks, supporting the proteomic evidence of a shift toward cytoplasmic localization with age. Similarly, LGALSL, a soluble β-galactoside-binding lectin present in both nuclear and cytoplasmic compartments (31), exhibited an N/C ratio of 1.287 ± 0.215 at 80 weeks, compared with 1.471 ± 0.323 at 35 weeks and 1.503 ± 0.371 at 11 weeks (Fig. 4d, e). Consistent with the proteomic data, this indicates a shift toward cytoplasmic localization during aging. Stratification of LGALSL-stained cells further demonstrated a progressive redistribution over time: cells with N/C ratios > 2 decreased markedly from 17.1% at 11 weeks to 11.9% at 35 weeks and 3.4% at 80 weeks, whereas cells with a more balanced distribution (N/C = 1.0–1.5) increased to 74.4% at 80 weeks (from 50.5% at both 11 and 35 weeks). In addition, the proportion of cells with predominantly cytoplasmic localization (N/C < 1), increased modestly from 3.8% at 11 weeks to 8.0% at 80 weeks (Fig. 4f).

SEC24C, a COPII coat cargo adaptor that mediates transport from the endoplasmic reticulum to the Golgi apparatus (32), exhibited a low N/C ratio at both 11 weeks (0.639 ± 0.168) and 35 weeks (0.657 ± 0.133), consistent with its mainly cytoplasmic localization in young podocytes (Fig. 5a, b). By 80 weeks, the N/C ratio increased to 0.735 ± 0.123, suggesting a relative shift toward nuclear enrichment. Analysis of N/C ratios revealed a gradual redistribution of signal over time. At 80 weeks, a small subset of cells (2.9%) exhibited an N/C ratio of 1.0–1.5, which was absent at 35 and 11 weeks (Fig. 5c). Most cells showed a shift toward a more balanced, yet slightly cytoplasm-dominant distribution: N/C ratios of 0.5–1.0 increased from 73.7% at 11 weeks to 85.3% at 35 weeks and reached 97.1% at 80 weeks. In contrast, cells with a strongly cytoplasmic signal (N/C < 0.5) significantly declined, disappearing completely at 80 weeks (0.0%) after being present in 14.7% of cells at 35 weeks and 26.3% at 11 weeks. These findings support the proteomic evidence of a progressive redistribution of SEC24C toward the nucleus during aging.

A similar pattern was observed for three mitochondrial proteins: NDUFA2 (Fig. 5d-f), ATP5PD (Supplemental Fig. 3a-c), and NDUFS6 (Supplemental Fig. 3d-f). All three showed low N/C ratios in the youngest group (0.678 ± 0.218, 0.499 ± 0.219, and 0.572 ± 0.161, respectively, at 11 weeks), which gradually increased with age. At 35 weeks, the ratios changed to 0.662 ± 0.234, 0.462 ± 0.173, and 0.611 ± 0.178, respectively, although ATP5PD did not show a significant increase at this stage. By 80 weeks, all three proteins showed significantly higher values (0.741 ± 0.277, 0.547 ± 0.242, and 0.656 ± 0.227, respectively). Across all mitochondrial protein stainings, we consistently observed a progressive enrichment of cells with N/C ratios in the 1.0–1.5 range (Fig. 5f; Supplemental Fig. 3c, f). In young podocytes, these proteins are mostly cytoplasmic, aligning with their mitochondrial localization. The gradual nuclear enrichment in aged cells reflects the proteomic finding that several mitochondrial proteins gain partial nuclear localization during aging (Fig. 3). One possible explanation is that declining mitochondrial function in older podocytes impairs efficient import of these proteins into mitochondria (33, 34), allowing a detectable nuclear pool to accumulate.

Together, these data indicate that aging alters the subcellular localization of cytoplasmic and nuclear proteins, consistent with impaired NPC barrier function. Because cellular processes most likely affected by this dysfunction include stress signaling and mitochondrial function, we examined both in aging mice. The transcriptional co-activator Yes-associated protein 1 (YAP1), a key effector of the Hippo pathway that controls growth and survival programs in response to metabolic and other stresses, exhibited significantly increased nuclear localization in podocytes from aged animals (Fig. 6a, b). N/C ratio analysis revealed a clear temporal shift: cells with an N/C ratio less than 1 markedly decreased from 38.8% at 11 weeks and 27.8% at 35 weeks to 15.9% at 80 weeks. In contrast, cells with a near-balanced distribution (N/C = 1.0–1.5) progressively increased, reaching 72.2% at 80 weeks compared to 51.8% at 11 weeks and 67.5% at 35 weeks. Additionally, cells with strongly nuclear-enriched localization (N/C >1.5) increased from 9.4% at 11 weeks and 4.7% at 35 weeks to 11.9% at 80 weeks (Fig. 6c). The nuclear accumulation of YAP1 in aged podocytes aligns with the reported upregulation of its downstream targets, including *Areg, Axl, Itgb2*, and *Myc*, which are increased by 5.7, 3.8, 5.5, and 2.7-fold, respectively, in aged versus young podocytes (35).

**Figure 6:**
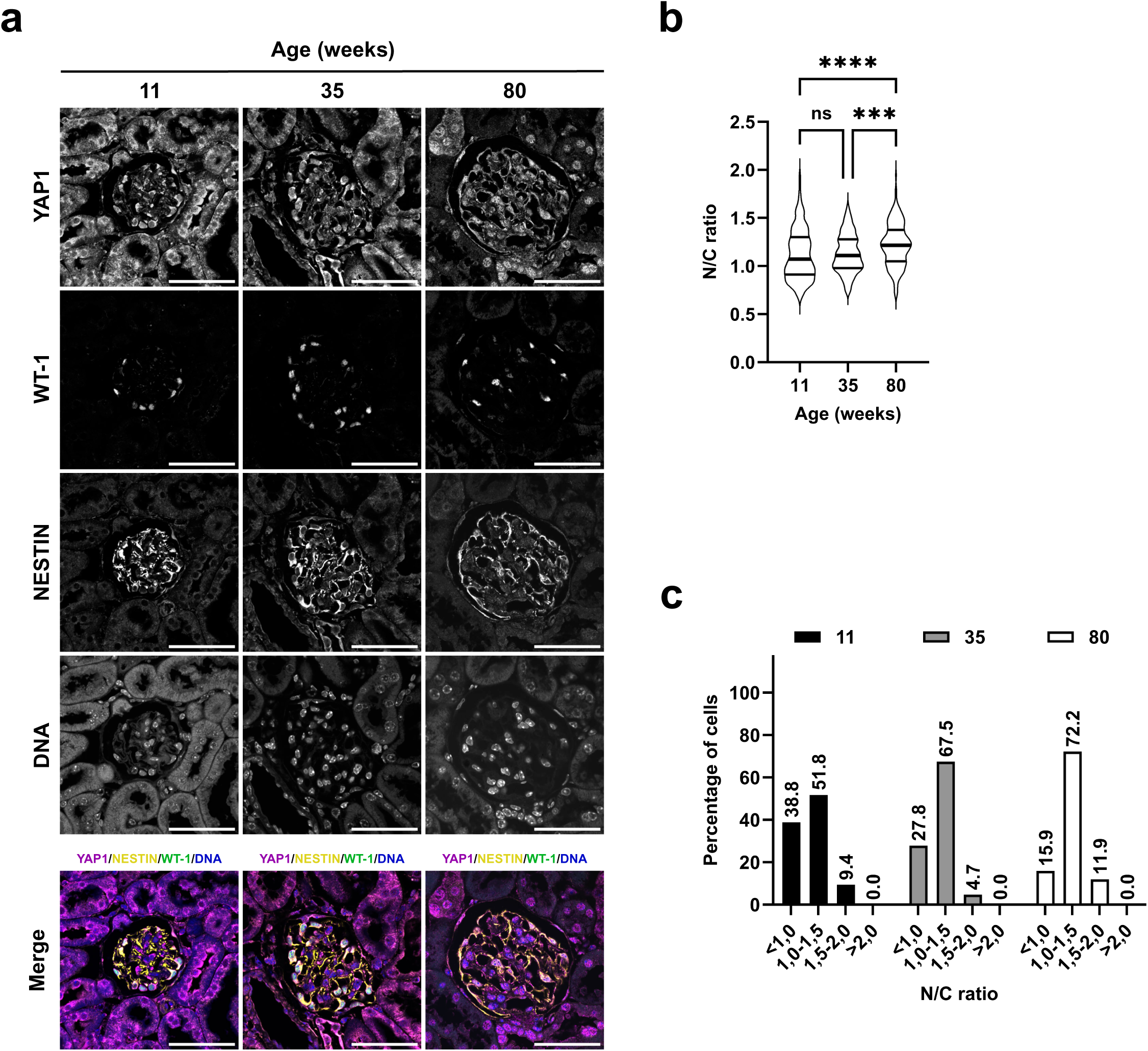
Aging promotes cytoplasmic-to-nuclear redistribution of YAP1 in podocytes. (a) Representative confocal images of kidney sections from 11, 35, and 80-week-old mice stained for YAP1 (magenta), with NESTIN marking the podocyte cytoplasm (yellow), WT-1 marking the podocyte nucleus (green), and DAPI labeling DNA (blue). Scale bar, 50 μm. (b) Nuclear-to-cytoplasmic ratio quantification of YAP1 staining intensity as in (**a**). Thick lines show the medians, thin lines indicate the 25% and 75% quartiles, with n = 174 analyzed podocytes at 11 weeks from 6 mice, n = 172 at 35 weeks from 6 mice, and n = 180 at 80 weeks from 6 mice. ns, not significant; ***P < 0.001; ****P< 0.0001. Kruskal-Wallis non-parametric test, one-way ANOVA, uncorrected Dunn’s test, multiple comparisons; 11 weeks vs. 35 weeks, P=00,3665, 11 weeks vs. 80 weeks, ****P < 0.0001, 35 weeks vs. 80 weeks, ***P=0.0003. (c) Distribution of podocytes into four nuclear-to-cytoplasmic ratio categories (NCs < 1, 1–1.5, 1.5–2, > 2), with percentages displayed above each column.

Conclusively, these quantitative imaging data corroborate both the direction and the magnitude of age dependent protein relocalization, providing independent validation of the nucleocytoplasmic ratio changes identified by proteomics.

### Podocytes from aged mice exhibit impaired mitochondrial function

To directly assess mitochondrial function in podocytes from young and aged mice, we measured mitochondrial membrane potential, a key indicator of mitochondrial activity and energy production. Podocytes were isolated from 11- and 80-week-old mice, and membrane potential was analyzed using tetramethylrhodamine ethyl ester (TMRE), a cell-permeable dye that accumulates in mitochondria in proportion to their membrane potential (36, 37). While 79.10% ± 2.46% of podocytes from young mice displayed high mitochondrial membrane potential, this proportion declined to 52.05% ± 15.34% in podocytes from aged mice, indicating impaired mitochondrial function (Fig. 7a). Decline in mitochondrial activity is often associated with pathological conditions and accompanied by increased superoxide production (38, 39). Consistently, staining with MitoSOX, a selective cell-permeable dye that detects mitochondrial superoxide, revealed positive signals in 52.45% ± 12.23% of podocytes from aged mice compared with 12.87% ± 1.97% in young mice (Fig. 7b). Together, these results indicate increased mitochondrial dysfunction in aged podocytes, characterized by reduced membrane potential and elevated superoxide levels.

**Figure 7:**
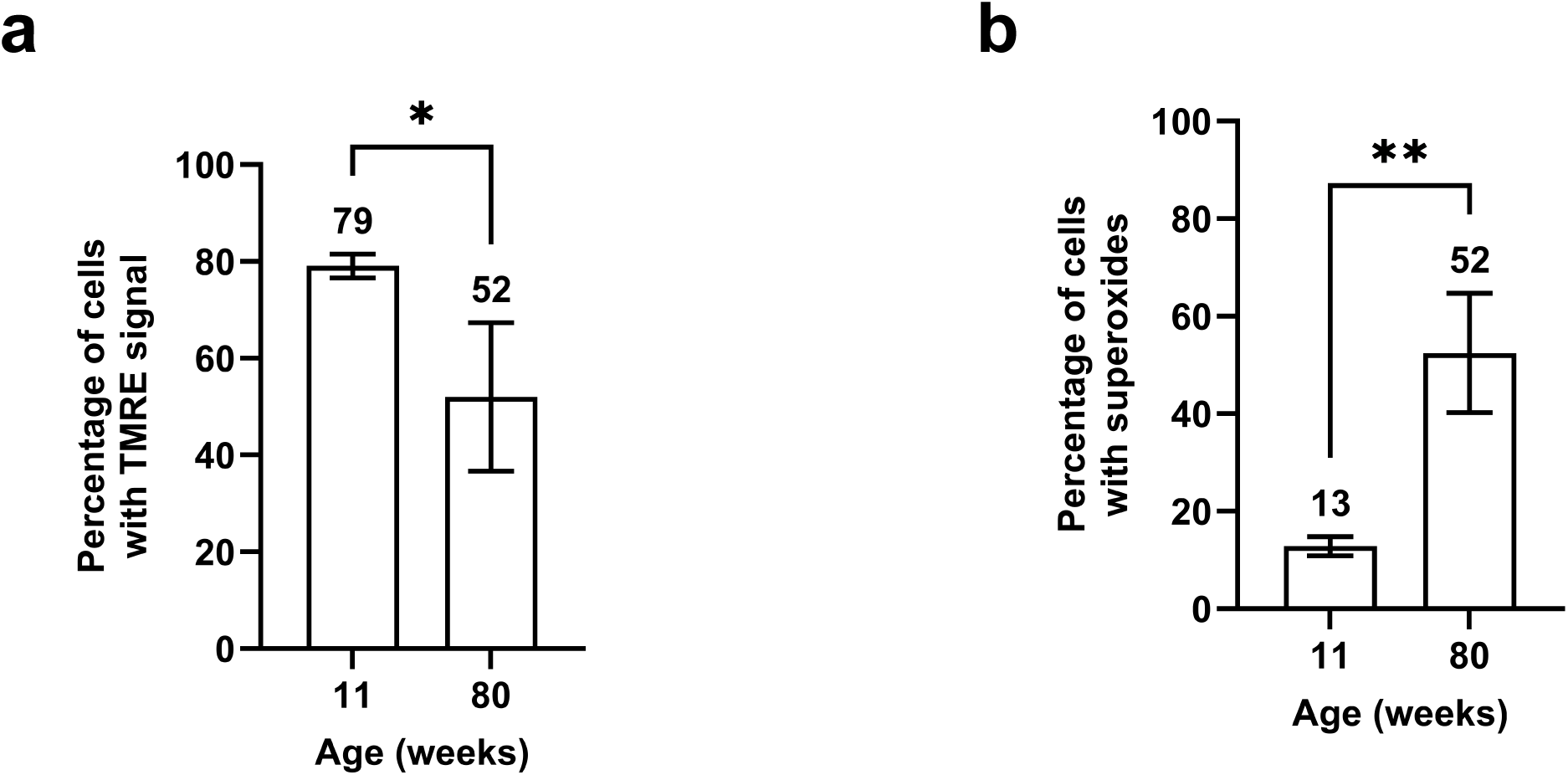
Aging causes mitochondrial membrane depolarization and increased superoxide production in podocytes. (a) Percentage of podocytes showing tetramethyl-rhodamine-ethyl-ester (TMRE) fluorescence from 11-week-old versus 80-week-old mice. Columns represent the mean ± standard deviation from independent biological replicates (n = 3 for 11 weeks, from 6 mice, and n = 2 for 80 weeks, from 4 mice). *P < 0.05. Unpaired experimental design, parametric test, unpaired t test, two-tailed. 11 weeks vs. 80 weeks, *P=0.0471. (b) Percentage of podocytes exhibiting MitoSOX fluorescence from 11-week-old and 80-week-old mice. mean ± standard deviation. n = 3 for 11 weeks, from 6 mice, and n = 2 for 80 weeks, from 4 mice. **P < 0.01. Unpaired experimental design, parametric test, unpaired t test, two-tailed. 11 weeks vs. 80 weeks, **P=0.0093.

### Mitochondrial dysfunction induces YAP1 nuclear localization in podocytes

Because aged podocytes display both increased stress signaling and mitochondrial dysfunction, we investigated whether these two phenomena are mechanistically connected. To this end, we used a well defined podocyte cell culture model (40, 41) that retains differentiation and expresses key podocyte markers, including PODOCIN, NEPHRIN, WT-1, and SYNAPTOPODIN (Supplemental Fig. 4a).

To test whether mitochondrial dysfunction directly activates the stress-responsive YAP1 pathway, we pharmacologically blocked the respiratory chain in cultured podocytes. Cells were treated with 10 µM antimycin A, a well-characterized Complex III inhibitor that increases mitochondrial superoxide production (42). MitoSOX fluorescence, measured as mean intensity, increased from 0.037 ± 0.002 in vehicle-treated control cells to 0.042 ± 0.004 following antimycin A exposure (Supplemental Fig. 4b, c), confirming the induction of mitochondrial oxidative stress. Quantitative immunofluorescence analysis revealed a clear shift of YAP1 toward the nucleus, with the N/C ratio increasing from 0.452 ± 0.160 in control cells to 0.625 ± 0.193 following antimycin A treatment (Fig. 8a, b).

Next, we induced mitochondrial dysfunction using 25 µM carbonyl cyanide m-chlorophenyl hydrazone (CCCP), a well-characterized mitochondrial uncoupler that collapses the proton gradient and eliminates the mitochondrial membrane potential (43). As a result, the mitochondrial membrane potential, assessed by mean TMRE fluorescence intensity, decreased from 0.323 ± 0.063 in untreated control cells to 0.040 ± 0.032 following CCCP treatment (Supplemental Fig. 4d, e). Along with the loss of membrane potential, YAP1 showed a significant shift toward the nucleus. Quantitative image analysis revealed that the N/C ratio of YAP1 increased to 0.595 ± 0.154 in CCCP treated cells, compared to 0.510 ± 0.129 in control cells (Fig. 8c, d).

Taken together, these results demonstrate that mitochondrial impairment, whether caused by respiratory chain inhibition or by collapse of the membrane potential, promotes nuclear translocation of YAP1 and most probable activates its downstream stress-responsive signaling pathway in podocytes.

### Mechanical stress induces mitochondrial dysfunction in podocytes

Mitochondrial dysfunction has been linked to mechanical stress in many tissues (44–46). For podocytes, the glomerular basement membrane is the primary source of intracellular mechanical cues (47, 48). To manipulate matrix stiffness, podocytes were cultured on substrates with defined elastic moduli, thereby varying the level of matrix rigidity (49). All substrates were coated with collagen IV and laminin to mimic the composition of the glomerular basement membrane. While 0.2 kPa and 64 kPa correspond to very soft and excessively stiff conditions, respectively, 8 kPa falls within the physiological range that supports healthy podocyte function (50). To assess the effects of matrix stiffness on cytoskeletal organization, we quantified stress fiber formation by measuring F-actin intensity. No significant differences in F-actin signal intensity were observed between 0.2 and 8 kPa (0.103 ± 0.053 and 0.112 ± 0.043, respectively, Supplemental Fig. 5a, b). In contrast, F-actin intensity increased in cells cultured at 64 kPa (0.170 ± 0.079), indicating that high matrix stiffness induces mechanical stress in podocytes. Consistently, vinculin-positive focal adhesion points were more prominent in cells cultured at 64 kPa compared with 0.2 and 8 kPa (Supplemental Fig. 5a). Furthermore, cells cultured at 64 kPa exhibited an increased N/C ratio for YAP1 (0.627 ± 0.290) compared to 0.2 kPa and 8 kPa (0.554 ± 0.282 and 0.488 ± 0.243, respectively; Supplemental Fig. 5c, d). Interestingly, the very soft matrix also caused a modest increase in nuclear YAP1 compared with physiological conditions, suggesting that both extremes of matrix stiffness can disrupt mechanotransduction and promote YAP1 nuclear localization (50).

To determine whether matrix rigidity and the associated mechanical stress directly impair mitochondrial function, we measured the mitochondrial membrane potential of podocytes cultured on substrates with three defined elasticities (Fig. 9a, b). Increasing substrate stiffness led to a gradual decline in TMRE fluorescence intensity (0.064 ± 0.007 at 0.2 kPa, 0.059 ± 0.005 at 8 kPa, and 0.056 ± 0.006 at 64 kPa). These data indicate that mechanical stress reduces mitochondrial membrane potential in this podocyte culture model. When mitochondrial superoxide production was assessed using MitoSOX, no detectable signal was observed under any stiffness condition (Supplemental Fig. 6), suggesting that loss of membrane potential occurs before, or independently of, increased reactive-oxygen species production.

**Figure 8:**
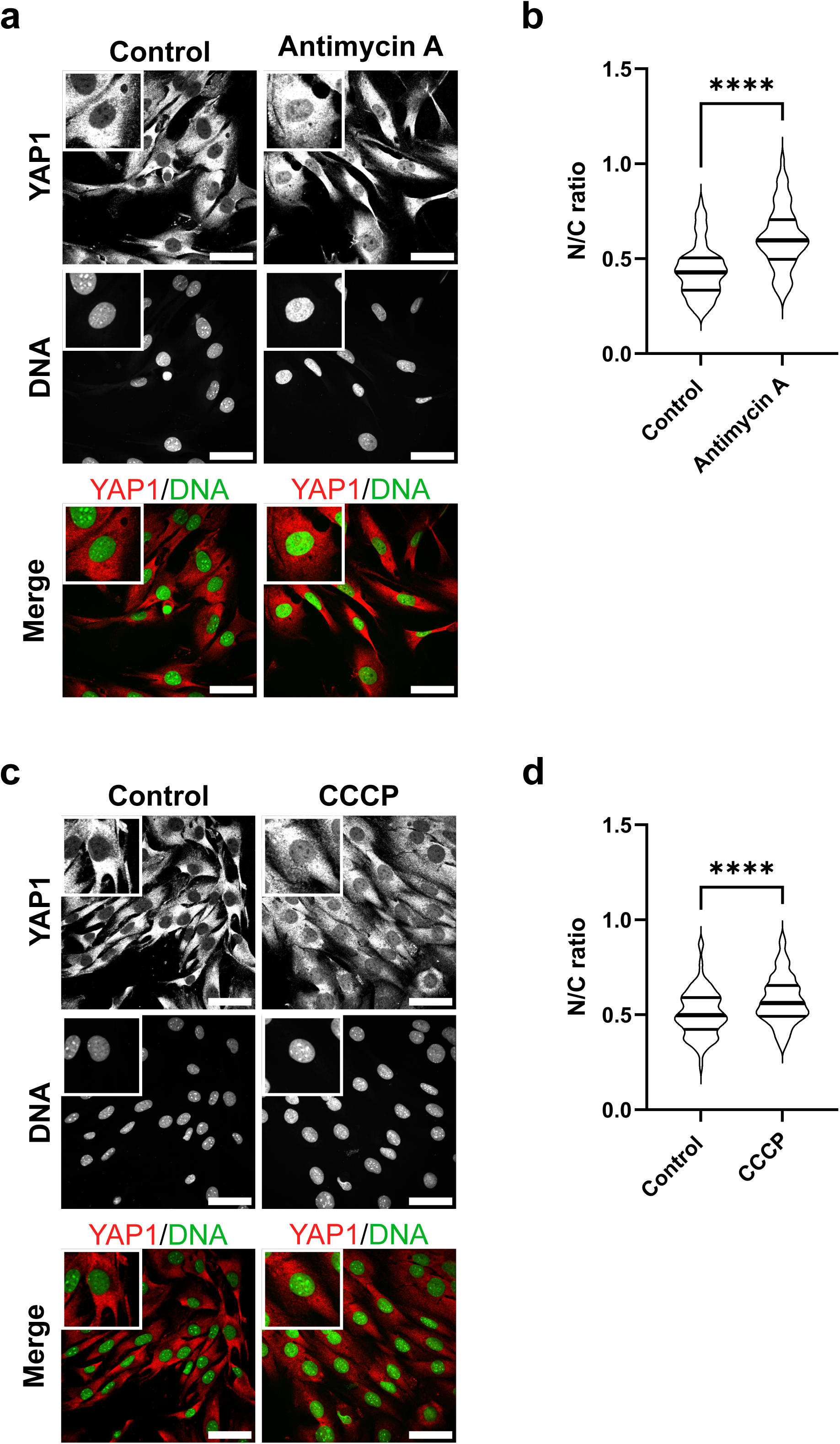
Mitochondrial stress induces YAP1 nuclear translocation in podocytes. (**a, c**) Representative confocal images of cultured podocytes, either control (DMSO) treated or exposed to 10 µM antimycin A (**a**) or 25 µM carbonyl cyanide m-chlorophenyl hydrazone (CCCP) (**c**) for 6 h, stained for YAP1 (red) and with DAPI (green). Scale bar, 50 μm. Insets show high-magnification views of individual cells. (**b**) Quantification of YAP1 nuclear-to-cytoplasmic ratio from three independent experiments, including a total of 171 podocytes in control conditions or after antimycin A treatment (n = 111). Thick lines indicate medians, and thin lines show the 25% and 75% quartiles. ****P<0.0001. Unpaired experimental design, nonparametric test, Mann-Whitney test, two-tailed. (**d**) Quantification of YAP1 nuclear-to-cytoplasmic ratio in control and CCCP-treated podocytes (control, n = 180; CCCP, n = 220), from three independent experiments. ****P< 0.0001. Unpaired experimental design, nonparametric test, Mann-Whitney test, two-tailed.

**Figure 9:**
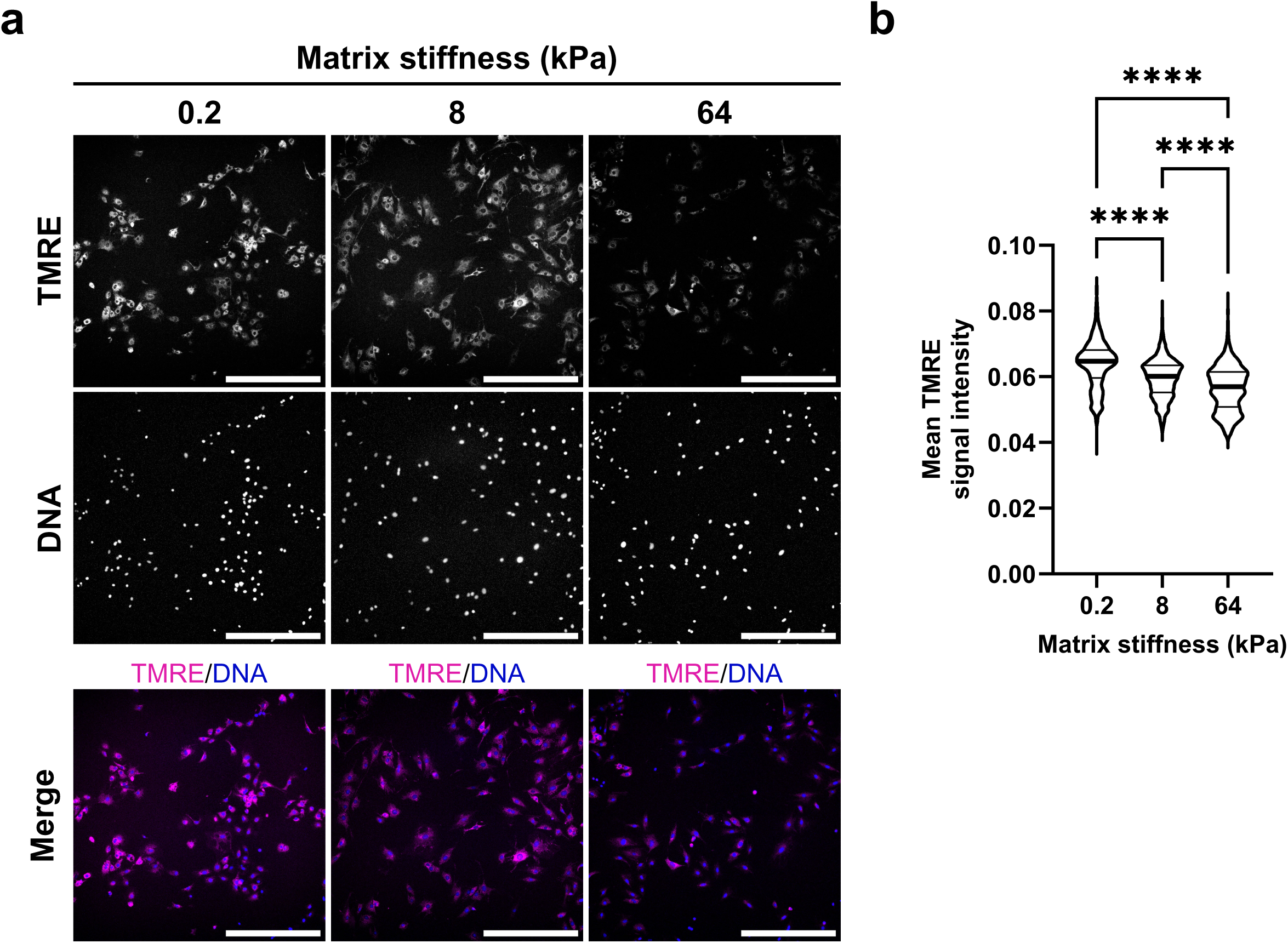
Matrix stiffness influences mitochondrial membrane potential in podocytes. (a) Representative confocal images of cultured podocytes grown on substrates of specific stiffness (0.2, 8, and 64 kPa) and stained with tetramethyl rhodamine ethyl ester (TMRE). Nuclei were visualized with Hoechst 33342. Scale bar, 500 μm. (b) Average TMRE fluorescence intensity measurements from samples as in (**a**), across three independent experiments. Thick lines indicate medians, while thin lines show the 25% and 75% quartiles. n = 1475 podocytes for 0.2 kPa, n = 1224 for 8 kPa, and n = 778 for 64 kPa. ****P< 0.0001. Kruskal-Wallis non-parametric test, one-way ANOVA, uncorrected Dunn’s test, multiple comparisons.

## Discussion

Podocytes are highly specialized epithelial cells that form a key component of the glomerular filtration barrier and are essential for lifelong kidney function. Because they are terminally differentiated and have virtually non-existent proliferative capacity, podocytes must maintain their structural integrity over decades. This remarkable longevity makes them particularly vulnerable to the cumulative cellular damage associated with aging (11, 51, 52). Here, we identify a previously unappreciated contributor in podocyte aging: the decline of NPC barrier function. Although the total number and density of NPCs remain largely unchanged with age, aged podocytes exhibit significantly increased NPC permeability. This loss of selective transport causes a widespread redistribution of proteins between the nucleus and cytoplasm, impacting factors involved in RNA processing, chromatin organization, and mitochondrial metabolism. These proteomic changes occur alongside mitochondrial dysfunction, increased production of reactive oxygen species, and enhanced nuclear localization of YAP1.

NPCs are essential for the compartmentalization of the nucleus and cytoplasm. In dividing vertebrate cells, these large structures are disassembled and reassembled during each mitosis (53). In contrast, postmitotic cells have a limited capacity to renew NPCs, and NPC dysfunction has been implicated as a contributing factor in cellular aging (54). In neurons, several nucleoporins that form the structural scaffold of the NPC, particularly NUP93 and NUP205, are remarkably long-lived proteins (19, 55). They exhibit very limited turnover, progressively accumulate oxidative damage, and consequently lead to progressive NPC leakiness and altered nuclear permeability. Related mechanisms have been associated with age-related neurodegenerative diseases, where impaired nucleocytoplasmic transport causes cellular dysfunction, mislocalization of RNA-binding proteins, and formation of pathogenic protein aggregates (56–58). Furthermore, significant metabolic remodeling linked to nuclear transport disruption has been observed in aging tissues (59, 60). Disruption of nucleocytoplasmic compartmentalization perturbs the nuclear localization of transcription factors, RNA-binding proteins, and chromatin modifiers, thereby altering gene expression (45, 46). In long-lived podocytes, loss of nuclear compartmentalization may gradually alter transcriptional programs that maintain the cytoskeleton and slit diaphragm, thereby weakening the glomerular filtration barrier. Similarly, transcriptional profiling of aged mice reveals significant downregulation of the slit diaphragm proteins nephrin and podocin, the transmembrane protein podocalyxin, cytoskeletal regulators such as synaptopodin and α-actinin-4, key components of the glomerular basement membrane including collagen type IV α4, elements of laminin-integrin signaling, the cell cycle inhibitor p57^Kip2, and the podocyte-specific transcription factors Lmx1b and WT 1, which are crucial for podocyte differentiation and function (35). In addition, aged podocytes show reduced transcriptional expression of VEGFA, a key endothelial survival factor required for maintenance of the glomerular filtration barrier (35).

A notable feature of our subcellular proteomics is the age-dependent mislocalization of mitochondrial proteins. Many mitochondrial proteins that accumulate in the nucleus of aged mice are smaller than the NPC diffusion cutoff of approximately 30 kDa, indicating that the age-related decline in NPC barrier integrity allows their passive entry. As mitochondrial protein-import efficiency declines with age (61, 62), these proteins probably persist longer in the cytoplasm, increasing the chance that they passively diffuse through NPCs. The age-related loss of NPC barrier integrity documented here may further exacerbate this effect. Additionally, weak or inactive NLS-like motifs may trigger nuclear import. Consistent with this, cNLS Mapper analysis identified such low-affinity NLS motifs in several mitochondrial proteins that become mislocalized to the nucleus in aged tissue (IARS2, ACSS3, PDHX, PDHB, SUOX, PDK2, ECSIT, GRSF1, MIEF1). ATFS1, a stress-activated transcription factor and the major regulator of the mitochondrial unfolded protein response, demonstrates how a protein normally imported into mitochondria can be redirected to the nucleus when mitochondrial import efficiency declines (63). Its inherent nuclear-localization signal becomes dominant, leading to nuclear accumulation under conditions of impaired mitochondrial import.

A decline in mitochondrial quality and activity is a hallmark of normal aging and is connected to many age-related diseases through multiple mechanisms (64, 65). For example, as oxidative phosphorylation becomes less efficient, ATP production decreases, thereby limiting the energy available for tissue maintenance and repair. Additionally, mitochondria produce increased levels of reactive oxygen species, which can damage proteins, lipids, and DNA. In line with these ideas, we observe mitochondrial dysfunction in aged podocytes, including loss of mitochondrial membrane potential and increased ROS levels. Mitochondrial injury has been linked to the development of kidney disease and is considered a hallmark of renal aging (66–68). Podocytes rely heavily on mitochondrial oxidative phosphorylation to meet the energy needs necessary to maintain their complex actin cytoskeleton and filtration barrier function (69, 70). Disruption of mitochondrial metabolism can therefore impair podocyte structure and survival (51, 71).

Our results show that aged mice have increased nuclear localization of YAP1 in podocytes. YAP1 is a key transcriptional regulator that integrates mechanical and metabolic signals to control cell adaptation and survival (72, 73). Mitochondrial dysfunction can affect YAP1 activity through changes in cellular energy status and cytoskeletal dynamics (74–76). Our finding that mitochondrial inhibitors promote YAP1 nuclear translocation in podocytes suggests that YAP1 may act as a sensor of metabolic stress in these cells, as previously observed in a breast cancer cell line (74). Activation of YAP1-driven transcription may initially serve as an adaptive response that supports cell survival under metabolic stress. However, ongoing YAP1 activation has been linked to fibrotic remodeling and glomerular disease (77–79), suggesting that sustained YAP1 activity may contribute to harmful changes in the aging kidney. Additionally, our results emphasize the influence of biomechanical factors on podocyte function. Podocytes reside in a dynamic mechanical environment, where forces generated by glomerular filtration continuously act on the filtration barrier (80). Mechanical stress affects cytoskeletal structure, mitochondrial function, and cellular metabolism across various cell types (81). The fact that abnormal matrix stiffness impairs mitochondrial function and promotes YAP1 nuclear localization suggests that mechanical alterations in the aging glomerulus may drive metabolic stress and transcriptional reprogramming in podocytes. Age-related shifts in extracellular matrix composition and glomerular structure (82, 83), combined with defects in nuclear permeability, may synergistically enhance cellular stress responses.

In summary, in a kidney-aging model, we observed a progressive decline in NPC function that coincides with increasing serum Cystatin C levels and the appearance of glomerular synechiae, early signs of podocyte injury that may ultimately lead to FSGS. Mutations in nucleoporin genes are known to cause a distinct subtype of FSGS that manifests as steroid-resistant nephrotic syndrome (SRNS) (20, 21, 84). In nucleoporin-linked SRNS, long-lived structural NPC components like NUP93 and NUP205 are frequently mutated; this loss of scaffold integrity weakens the NPC barrier and impairs overall nucleocytoplasmic transport (20, 21, 85). This disruption is believed to change gene expression, stress-response pathways, and glomerular development. Notably, a rat model of FSGS showed that pharmacological inhibition of nuclear transport, which restores the balance of nuclear-to-cytoplasmic protein distribution, significantly improved kidney function (29). It will be important to explore how, beyond the common defect in nucleocytoplasmic compartmentalization, nucleoporin-associated SRNS phenotypically overlaps with the podocyte aging phenotype described here. Understanding this connection could uncover shared molecular pathways that promote podocyte degeneration and suggest new therapeutic strategies for both age-related and genetic forms of FSGS.

## Methods

### Mouse Experimental Model

Podocyte-specific reporter mice (Pod-rtTA/LC1/R26R/H2B-eGFP; inbred for more than 20 generations with over 90% Sv129 genetic background; hereafter referred to as Pod-rtTA mice) were housed at the animal facility of Aachen University Hospital under specific pathogen-free conditions, with a 12-hour light/dark cycle and controlled temperature and humidity. The animals had *ad libitum* access to standard chow and water. All animal experiments and procedures adhered to institutional and national guidelines and were approved by the appropriate governmental authority (LAVE NRW; approval no. 81-02.04.2021.A471).

These mice were used to identify and isolate podocytes. In this doxycycline-inducible system, the podocyte-specific NPHS2 (podocin) promoter drives the expression of the reverse tetracycline-controlled transactivator (rtTA), which induces the expression of a histone H2B-eGFP fusion protein, resulting in nuclear eGFP labeling of podocytes (40, 86). No phenotypic abnormalities were observed in mice of the respective age groups following induction (see protocol below). For podocyte isolation, glomeruli were isolated, and primary podocytes were identified based on GFP fluorescence and subsequently purified by fluorescence-activated cell sorting (FACS; see below).

### Podocyte isolation, cell fractionation

Histon H2B-eGFP fusion protein expression to label podocytes was induced in mice 14 days prior to sacrifice by administering drinking water supplemented with 2% doxycycline and 5% sucrose. The drinking water was replaced every other day to maintain doxycycline activity.

Podocytes were isolated from mouse kidneys as previously described (41), with slight modifications. Briefly, kidneys were harvested, decapsulated, minced, and enzymatically digested in HBSS supplemented with collagenase Type 2 (LS004174, Worthington, Lakewood, NJ, USA), Protease (P5147, Merk, Germany), and DNase I (9003-98-9; PanReac AppliChem GmbH, Darmstadt, Germany). For the initial digestion, 1.5 ml of digestion buffer per kidney was used, and samples were incubated for a total of 15 minutes with gentle disruption by pipetting up and down every 5 minutes. Glomeruli were enriched by differential sieving through 100 µm and 40 µm nylon mesh cell strainers (Corning Inc., Corning, NY, USA). To generate single cells, isolated glomeruli were resuspended in 2 ml of the digestion buffer (see above) per mouse and incubated on a horizontal shaker at 1400 rpm in 5-minute intervals until single podocytes, identified by fluorescence microscopy by nuclear histone-eGFP expression, were obtained. Between incubation intervals, glomeruli were mechanically disrupted by repeated pipetting, vortexing, and passage through a syringe to ensure complete detachment of podocytes. Cell suspensions were washed, resuspended in ice-cold FACS buffer (HBSS + 0.5% BSA + 2 mM EDTA), and filtered through a 40 µm mesh (Corning Inc.). Subsequently, GFP-positive podocytes were sorted at 37 °C using a Becton Dickinson FACSAria™ fusion flow cytometer (BD Biosciences, San José, CA, USA) with identical acquisition settings for all samples. Forward and side scatter gating was applied to exclude debris and cell aggregates. Isolated podocytes were further used for cell fractionation, evaluation of NPC permeability by dextran, or assessing mitochondrial integrity as described below.

Cell fractionation was carried out by pelleting and washing freshly isolated podocytes in fractionation buffer (10 mM HEPES (pH 7.9), 10 mM KCl, 1.5 mM MgCl_2_, supplemented with 0.5 µl/ml DTT and protease inhibitor mix (IP100, Thermo Fisher, MA, USA)). Cells were resuspended in a minimal volume of fractionation buffer containing 0.1% NP-40 and frozen at -80°C. Frozen podocytes were thawed on ice for 5 minutes and centrifuged at 2,600 rpm for 10 minutes at 4 °C in a tabletop centrifuge to pellet nuclei. The resulting supernatant was collected as the cytoplasmic fraction. The nuclear pellet was washed three times with ice-cold fractionation buffer, then resuspended in a minimal volume of 20 µl in nuclear buffer (50 mM HEPES (pH 7.9), 150 mM NaCl, 0.2 mM EGTA, 0.2 mM EDTA, 0.5% C_24_H_39_NaO_4_) and kept on ice for 20 minutes with intermittent vortexing. The nuclear fraction was separated by centrifugation at 13,000 rpm for 10 minutes at 4 °C to pellet membrane components, and the supernatant was collected as the nuclear fraction.

Protein concentrations in the collected cytoplasmic and nuclear fractions were determined using the NanoOrange™ Protein Quantitation Kit according to the manufacturer’s protocol. Fraction purity was verified by Western blotting (see below). Membranes were probed with antibodies against histone H3 (rabbit monoclonal, Cell Signaling Technology #4499; nuclear marker) and β-actin (mouse monoclonal, Sigma-Aldrich #A1978; cytoplasmic marker) or vinculin (clone hVIN-1, mouse monoclonal, Sigma-Aldrich #V9131; cytoplasmic marker). Secondary antibodies were anti-mouse IgG (sheep polyclonal, HRP-conjugated, Abcam #ab6808) and anti-rabbit IgG (goat polyclonal, HRP-conjugated, Merck #A6154). Histone H3 was detected exclusively in nuclear fractions, whereas β-actin and vinculin were restricted to cytoplasmic fractions, confirming negligible cross-contamination and sufficient purity for subsequent mass spectrometric analysis.

### Mass spectrometry

Protein samples were subjected to the Single-Pot Solid-Phase-enhanced Sample Preparation (SP3) protocol (87) conducted on the KingFisher Apex™ platform (Thermo Fisher Scientific). For digestion, trypsin was used in a 1:20 ratio (protease: protein) in 50 mM N-2-hydroxyethylpiperazine-N-2-ethanesulfonic acid (HEPES) supplemented with 5 mM Tris(2-carboxyethyl) phosphine hydrochloride (TCEP) and 20 mM 2-chloroacetamide (CAA). Digestion was carried out for 5 hours at 37°C.

Up to 10 µg of peptides were labeled using TMTpro™ 16plex reagent as previously described (88). Briefly, 0.5 mg of TMT reagent was dissolved in 45 µL of 100% acetonitrile. Subsequently, 4 µL of this solution was added to each peptide sample, and the samples were incubated at room temperature for 1 hour. The labeling reaction was quenched by adding 4 µL of a 5% aqueous hydroxylamine solution and incubating for an additional 15 minutes at room temperature. Labeled samples were then combined for multiplexing, desalted using an Oasis® HLB µElution Plate (Waters) according to the manufacturer’s instructions, and dried by vacuum centrifugation.

Offline high-pH reversed-phase fractionation (89) was carried out using an Agilent 1200 Infinity high-performance liquid chromatography (HPLC) system, equipped with a Gemini C18 analytical column (3 μm particle size, 110 Å pore size, dimensions 100 x 1.0 mm, Phenomenex) and a Gemini C18 SecurityGuard pre-column cartridge (4 x 2.0 mm, Phenomenex). The mobile phases consisted of 20 mM ammonium formate adjusted to pH 10.0 (Buffer A) and 100% acetonitrile (Buffer B). The peptides were separated at a flow rate of 0.1 mL/min using the following linear gradient: 100% Buffer A for 2 minutes, ramping to 35% Buffer B over 59 minutes, increasing rapidly to 85% Buffer B within 1 minute, and holding at 85% Buffer B for an additional 15 minutes. Subsequently, the column was returned to 100% Buffer A and re-equilibrated for 13 minutes. During the LC separation, 48 fractions were collected. These were pooled into twelve fractions by combining every twelfth fraction. The pooled fractions were then dried using vacuum centrifugation.

An UltiMate 3000 RSLCnano LC system (Thermo Fisher Scientific) equipped with a trapping cartridge (µ-Precolumn C18 PepMap™ 100, 300 µm i.d. × 5 mm, 5 µm particle size, 100 Å pore size; Thermo Fisher Scientific) and an analytical column (nanoEase™ M/Z HSS T3, 75 µm i.d. × 250 mm, 1.8 µm particle size, 100 Å pore size; Waters) was used. Samples were trapped at a constant flow rate of 30 µL/min using 0.05% trifluoroacetic acid (TFA) in water for 6 minutes. After switching in-line with the analytical column, which was pre-equilibrated with solvent A (3% dimethyl sulfoxide [DMSO], 0.1% formic acid in water), the peptides were eluted at a constant flow rate of 0.3 µL/min using a gradient of increasing solvent B concentration (3% DMSO, 0.1% formic acid in acetonitrile).

Peptides were introduced into an Orbitrap Fusion™ Lumos™ Tribrid™ mass spectrometer (Thermo Fisher Scientific) *via* a Pico-Tip emitter (360 µm OD × 20 µm ID; 10 µm tip, CoAnn Technologies) using an applied spray voltage of 2.2 kV. The capillary temperature was maintained at 275 °C. Full MS scans were acquired in profile mode over an m/z range of 375–1,500, with a resolution of 120,000 at m/z 200 in the Orbitrap. The maximum injection time was set to 50 ms, and the AGC target limit was set to ‘standard’. The instrument was operated in data-dependent acquisition (DDA) mode, with MS/MS scans acquired in the Orbitrap at a resolution of 30,000. The maximum injection time was set to 94 ms, with an AGC target of 200%. Fragmentation was performed using higher-energy collisional dissociation (HCD) with a normalized collision energy of 34%, and MS2 spectra were acquired in profile mode. The quadrupole isolation window was set to 0.7 m/z, and dynamic exclusion was enabled with a duration of 60 seconds. Only precursor ions with charge states 2–7 were selected for fragmentation.

### Database search and proteomics data analysis

Raw files were converted to mzML format using MSConvert from ProteoWizard, with peak picking, 64-bit encoding, zlib compression, and filtering for the 1000 most intense peaks. Files were then searched using MSFragger (v4.0) in FragPipe (21.1) against the mus musculus FASTA database UP000000589 (C57BL, ID10090, 21,968 entries, October 2022), which contains common contaminants and reversed sequences. The following modifications were included in the search parameters: Carbamidomethylation (C, 57.0215), TMTpro (K, 304.2072) as fixed modifications; Oxidation (M, 15.9949), Acetylation (protein N-terminus, 42.0106), TMTpro (peptide N-terminus, 304.2072) as variable modifications. For the full scan (MS1), a mass error tolerance of 20 PPM was set, and for MS/MS (MS2) spectra, 20 PPM. Protein digestion used trypsin as the protease, allowing a maximum of two missed cleavages and requiring a minimum peptide length of seven amino acids. The false discovery rate was set to 0.01 at both peptide and protein levels. The standard settings of the FragPipe workflow ‘TMT16’ were used, with the following modifications: msfragger.misc.fragger.enzyme-dropdown-1: trypsin, msfragger.misc.fragger.precursor-charge-hi: 6, msfragger.search enzyme name 1: trypsin, msfragger.search enzyme nocut 1: P, msfragger.use topN peaks: 300, tmtintegrator.allow unlabeled: true, tmtintegrator.extraction tool: Philosopher.

For proteomics data analysis, the raw output files from FragPipe ((90), protein.tsv files) were processed in the R programming environment (ISBN 3-900051-07-0). Initial data processing included filtering out contaminants and reverse proteins. Only proteins quantified with at least 2 razor peptides (Razor.Peptides >= 2) were considered for further analysis. 3594 proteins passed the quality control filters. Abundance values (’channel’ columns) were log2-transformed and assembled into a Bioconductor ExpressionSet object for downstream analysis. Infinite values resulting from log2 transformation of zero values were set to NA. Log2-transformed raw TMT reporter ion intensities were normalized using the ‘normalizeVSN’ function of the limma package (VSN - variance stabilization normalization - (91)). This normalization was performed across all TMT channels, including cytosolic and nuclear fractions, to correct for global differences in reporter ion intensity distributions between samples. Nuclear-to-cytosolic ratios were then calculated for each protein by dividing the normalized nuclear abundance by the median normalized abundance of the corresponding cytosolic condition at the same age group, e.g. nuclear 11 weeks / cytosol 11 weeks, nuclear 35 weeks / cytosol 35 weeks, and nuclear 80 weeks / cytosol 80 weeks. These ratios were used for clustering, heatmap visualization, and downstream comparison of age-dependent changes in subcellular distribution. Because normalization aligns the global intensity distributions of nuclear and cytosolic samples, the resulting N/C ratios reflect relative protein-specific deviations from the global distribution rather than absolute differences in total protein abundance between fractions. Differential expression analysis was performed using the moderated t-test provided by the limma package (92). The model accounted for replicate information by including it as a factor in the design matrix passed to the ‘lmFit’ function. Proteins were annotated as hits if they had a false discovery rate (FDR) below 0.05 and an absolute fold change greater than 1.5. Proteins were considered candidates if they had an FDR below 0.2 and an absolute fold change greater than 1.5.

### Cell Culture

For cell culture experiments, primary mouse podocytes were isolated from the kidneys of Pod-rtTA mice as previously described (41). Following isolation, cells were cultured in RPMI 1640 medium (#21875091, Gibco, Thermo Fisher Scientific, Waltham, MA, USA) supplemented with 10% (v/v) heat-inactivated fetal bovine serum (FBS; #17593595, Gibco, Thermo Fisher Scientific, Waltham, MA, USA) and 1% (v/v) penicillin/streptomycin (#15140122, Gibco, Thermo Fisher Scientific, Waltham, MA, USA) at 37°C in a humidified atmosphere containing 5% CO_2_. The cells were regularly tested and confirmed to be free of mycoplasma.

For the analysis of podocyte mechanobiological properties, cytosoft® 6-well plates with defined substrate stiffnesses (0.2, 8, and 64 kPa; Sigma-Aldrich) were coated with collagen IV (3 µg/cm²; #CLS354233, Corning Inc., Corning, NY, USA) and laminin (5 µg/cm²; #CLS354232, Corning Inc., Corning, NY, USA). Collagen IV and laminin were diluted in cold phosphate-buffered saline (PBS) to the desired final concentrations, and the coating solution was evenly applied to each well. Plates were incubated for 1 hour at room temperature under sterile conditions. After incubation, the coating solution was removed, and wells were washed with cold PBS. Plates were either used immediately or sealed and stored at 4 °C until further use. 1.0 × 10 podocytes were then seeded onto the coated substrates and maintained at 37 °C in a humidified incubator with 5% CO.

### Mitochondrial membrane potential and mitochondrial superoxide assays

To measure mitochondrial membrane potential and superoxide production, primary mouse podocytes were cultured under standard conditions as described above. Mitochondrial membrane potential was assessed using tetramethylrhodamine ethyl ester (TMRE; #Cay21437, Invitrogen), and mitochondrial superoxide production was measured using MitoSOX Red (#M36007, Invitrogen). Both parameters were analyzed using live cell imaging.

For live imaging of TMRE staining, podocytes in culture were incubated with TMRE diluted in pre-warmed full cell culture medium (final concentration 400 nM) for 30 minutes at 37°C, protected from light. As a positive control for mitochondrial depolarization, cells were treated with carbonyl cyanide m-chlorophenyl hydrazone (CCCP; 25 µM; # CS-7660, Chemscene) for 6 hours before TMRE staining. For live imaging of MitoSOX staining, cells were incubated with MitoSOX Red (final concentration 500 nM) in serum-free medium for 30 minutes at 37°C, protected from light. Antimycin A (10 µM; J63522-LB0, Thermo Fisher Scientific) was used for 6 hours before MitoSOX staining as a positive control to induce mitochondrial superoxide production. After incubation, cells were gently washed with pre-warmed PBS and immediately imaged in phenol red–free medium. Fluorescence images were captured using a spinning-disk Ti2-E confocal microscope (Nikon, Melville, NY, USA) equipped with a Plan-Apochromat ×10 NA 0.5 objective, an environmental control system (UNO-T-H-CO; Okölab, Ottaviano, Italy), a SpectraX LED light engine with filter sets for DAPI and Cy5 (Lumencor, Beaverton, OR, USA), and a Digital Sight 10 CCD camera (Nikon). Image acquisition was performed with NIS-Elements software (Nikon). Image processing and quantitative analysis were performed using ImageJ and CellProfiler, with identical acquisition and processing parameters applied across all experimental groups.

To measure mitochondrial membrane potential and superoxide production in isolated podocytes, freshly isolated podocytes from 11- and 80-week-old mice were stained in pre-warmed FACS buffer (HBSS+ 2% BSA) at 37 °C. Throughout all staining steps, cells were incubated on a horizontal shaker at 300 rpm to ensure gentle and uniform suspension. To assess mitochondrial membrane potential, cells were incubated with TMRE (final concentration: 400 nM) for 30 minutes at 37 °C in the dark. To measure mitochondrial superoxide, cells were incubated with MitoSOX Red (final concentration 500 nM) for 30 minutes at 37 °C in the dark. After incubation, samples were diluted fivefold in FACS buffer to reduce background signal and immediately analyzed on a Becton Dickinson FACSCanto^TM^ II flow cytometer (BD Biosciences, San José, CA, USA), and data were analyzed using BD FACSDiva software (BD Biosciences, San José, CA, USA). Debris and dead cells were gated out based on forward and side scatter. TMRE and MitoSOX fluorescence were presented as the percentage of cells.

### Western blotting

Proteins were denatured at 95 °C for 5 min, adjusted to equal concentrations, mixed with loading buffer (4:1), and separated on 10% SDS-PAGE at 120 V. Proteins were transferred to PVDF membranes (Merck Millipore, Massachusetts) at 100 V for 1 h, blocked with 2% BSA in TTBS for 1 h at room temperature, and incubated with primary antibodies for 1 h at room temperature or overnight at 4 °C. After five washes in TBS with 0.1% Tween-20, membranes were incubated with HRP-conjugated secondary antibodies for 1 h at room temperature. Signals were detected using Lumi-Light chemiluminescent substrate (Roche, distributed by Sigma-Aldrich/Merck) following the manufacturer’s instructions.

### Nuclear permeability assays

Freshly isolated podocytes were pelleted and washed in fractionation buffer (10 mM HEPES, 10 mM KCl, 1.5 mM MgCl_2_, supplemented with 0.5 µl/ml DTT and protease inhibitor mix (IP100, Thermo Fisher, MA, USA)). To permeabilize the cell membrane, the pellet was resuspended in fractionation buffer containing 0.2% NP-40 and incubated on ice for 15 minutes, with intermittent vortexing every 5 minutes. The permeabilized cells were then washed and resuspended in a minimal volume of fractionation buffer.

To assess nuclear permeability, cells were mixed with dextrans of 10-, 40-, or 70-kDa at a 1:1 ratio (25 mg/ml in water, D1828, #D1829, and #D1830, respectively, Thermo Fisher Scientific) and incubated on ice for 10 minutes. The cell-dextran mixtures were then transferred to microscope slides and analyzed *via* fluorescence microscopy.

### Serum collection and analyses

Blood samples were collected *via* retro-orbital bleeding, serum was then isolated, and serum Cystatin C levels were measured using an ELISA kit (R&D Systems, Minneapolis, MN, USA), following the manufacturer’s instructions.

### Podocyte counting

Kidney tissues were collected. Approximately 1 mm-thick tissue sections were dissected and immediately immersed in 4% paraformaldehyde (PFA) for 24 hours, then washed in phosphate-buffered saline (PBS) for 12 hours to remove residual fixative. Tissue dehydration involved incubating the samples in fresh absolute ethanol for two cycles of 30 minutes each. All incubation steps were performed on a shaker at room temperature. After dehydration, tissues were transferred into pure ethyl cinnamate for optical clearing until they became visibly transparent at room temperature.

For podocyte counting, three-dimensional imaging was performed using an Olympus FV1000MPE two-photon laser scanning microscope equipped with a 25x water immersion objective (numerical aperture: 1.05) and a correction collar for refractive index adjustment. An excitation wavelength of 820 nm was used to detect the eGFP signal. Image processing and analysis were carried out using Imaris software, specifically the “Spots” function. For each experimental group, kidney tissues from 5-8 animals were analyzed, and 8-10 glomeruli per animal were evaluated.

### Histochemistry, and Immunofluorescence

Kidney tissue samples were fixed in methyl-Carnoy’s solution (MethaCarn; 60% methanol, 30% chloroform, 10% glacial acetic acid), 3% paraformaldehyde (PFA), or 4% formaldehyde, and then embedded in paraffin. Before staining, sections were deparaffinized in xylene and rehydrated through a graded ethanol series.

To quantify glomerular synechiae, a hallmark of early FSGS, 1 µm-thick sections of MethaCarn-fixed kidney tissue from all age groups (n=7 for the 11-week group, n=8 for the 35- and 80-week groups) were stained with periodic acid-Schiff’s reagent (PAS) and counterstained with hematoxylin. All glomeruli in each PAS-stained tissue section were examined at 400X magnification using a light microscope (Olympus BX41), and the percentage of glomeruli exhibiting synechiae was determined for each age group by an investigator blinded to group allocation.

For immunofluorescence, formalin- and PFA-fixed sections were subjected to antigen retrieval by microwaving in a citrate-based antigen-unmasking solution at 600 W for 3 cycles of 5 minutes each. Sections were subsequently blocked with 3% BSA in PBS for 1 hour in a humidified chamber, and incubated with primary antibodies for 1 hour at room temperature or overnight at 4 °C, followed by incubation with fluorescence-labeled secondary antibodies for 45 minutes. Sections were incubated with DAPI (1 µg/ml, Roche Diagnostics GmbH, Mannheim, Germany) for 10 minutes, mounted with Immu-Mount, and covered with #1.5H high-precision glass coverslips (170 µm thickness).

To quantify nuclear-to-cytoplasmic signal ratios in kidney sections, tissues were immunolabeled with the respective primary antibodies and counterstained with WT-1 and NESTIN as podocyte-specific nuclear and cytoplasmic markers, respectively. Sections were analyzed at 630-fold magnification using a Zeiss LSM 710 inverted confocal laser scanning microscope (Zeiss, Oberkochen, Germany) equipped with a Plan Apochromat ×63/1.4 immersion oil objective. For quantifying nuclear-to-cytoplasmic signal ratios of dextran in permeabilized podocyte cells, images were taken at 200-fold magnification with a BZ9000 fluorescence microscope (Keyence, Neu-Isenburg, Germany) fitted with a Plan Apochromat ×20/0.75 air objective. Nuclear-to-cytoplasmic ratios were calculated using a custom ImageJ pipeline for image pre-processing, followed by a custom CellProfiler pipeline for downstream analysis, as previously described (29). Immunofluorescence staining of NDUFA2, NDUFS6, and ATP5PD was performed on 1 µm-thick formalin-fixed sections using a triple-staining protocol with NESTIN and WT-1. IF staining of LGALSL, SERPINB6, SEC24C, and YAP1 was conducted on 1 µm-thick PFA-fixed sections with a triple-staining approach including NESTIN and WT-1, to determine the podocyte-specific nuclear-to-cytoplasmic signal intensity ratio. All antibodies used were tested and validated for subsequent experiments, with staining patterns consistent with previous reports.

To quantify NPCs in podocytes by IF, 3 µm-thick PFA-fixed sections were stained with WT-1 and mAB414 and analyzed using stimulated emission depletion (STED) microscopy. Sections stained with mAB414 and anti-WT-1 antibodies were imaged using an inverted Leica SP8 Tau-STED3X microscope (Leica Biosystems Nussloch GmbH, Nußloch, Germany) equipped with a Leica HC PL APO 93× NA 1.30 glycerol-immersion STED objective and appropriate filter settings. Image processing and analysis were performed using ImageJ and Imaris. For each cell, a z-stack of four optical sections was acquired, and a mean-intensity projection was created to generate a single representative image. Nuclear WT-1 signals were segmented using the Imaris “Surface” module with automatic thresholding. Nuclear pore complex (NPC) signals labeled with mAB414 were identified with the Imaris “Spots” module, using automatic thresholding and a defined spot diameter of 100 nm. The number of NPCs and the distances to the three nearest neighboring NPCs were quantified with Imaris.

Details of the primary and secondary antibodies used are provided in Supplemental table 1.

### Statistical Analyses

Results are shown as mean ± standard deviation (SD) or median with 25% and 75% quartiles as indicated and where appropriate. Statistical analyses were conducted using GraphPad Prism (version 10.4.1; GraphPad Software Inc., San Diego, CA, USA). Outliers were identified and removed using the ROUT method with a false discovery rate (FDR) of 10% for each dataset. Only cleaned datasets were used in subsequent analyses. Data normality was tested with the Shapiro-Wilk test. Comparisons between two groups were made using a two-tailed Student’s t test for normally distributed data (α = 0.05) and the Mann-Whitney U test for non-normal data (α = 0.05). When comparing more than two groups, one-way analysis of variance (ANOVA) followed by Fisher’s least significant difference (LSD) post hoc test was used for parametric data, while the Kruskal-Wallis test was applied for nonparametric data.

## Acknowledgements

We acknowledge the excellent technical assistance of Mihir Patel, and Susanne Scholz. This study was supported by the Immunohistochemistry, Two-Photon Imaging and Confocal Microscopy facilities, which are core facilities of the Interdisciplinary Center for Clinical Research (IZKF) Aachen, at the Faculty of Medicine, RWTH Aachen University. This work was further supported by the Flow Cytometry Facility, a core facility of the Interdisciplinary Center for Clinical Research (IZKF) Aachen within the Faculty of Medicine at RWTH Aachen University and by the German Research Foundation (DFG), project ID 439895892. We further acknowledge the Core Facility for 3D Super-Resolution (3DSR) at RWTH Aachen for imaging assistance.

## Author contributions

M.H., Y.G., T.O., W.A.: Conceptualization; M.H., Y.G., F.S.: formal analysis; M.H., Y.G., G.V., T.S., U.R., I.M.: investigation; E.S., R.J., U.R., R.K.: resources; M.H., T.O, W.A.: writing - original draft preparation; M.H., Y.G., R.J., T.O., W.A.: writing - review and editing; M.H., Y.G.: visualization; T.O., W.A.: supervision and project administration; M.H., R.K., T.O., W.A.: funding acquisition. All authors have read and agreed to the submitted version of the manuscript.

---

**Supplemental Figure 1:**
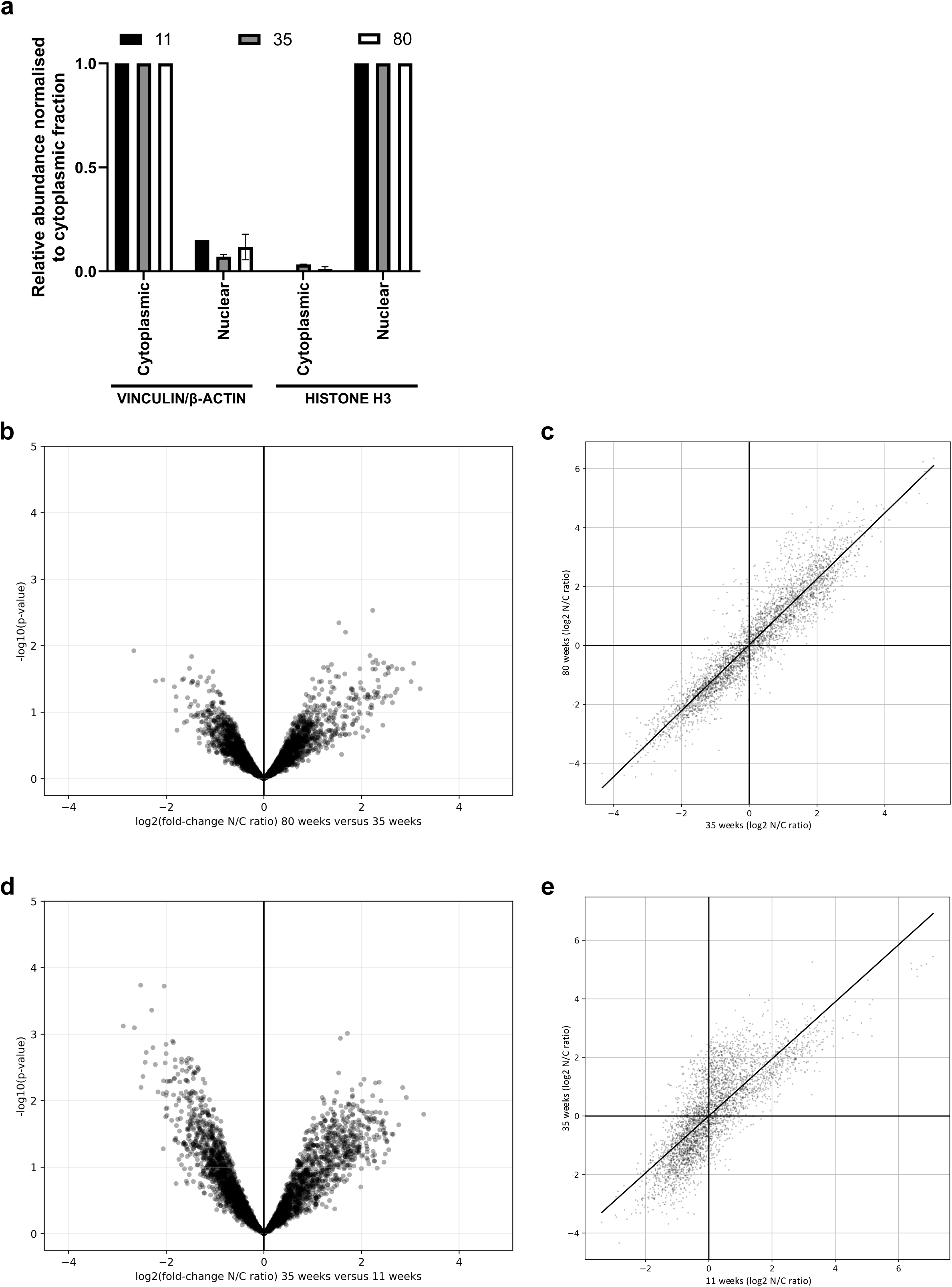
Age-dependent changes in subcellular protein distribution in podocytes. (a) Quantification of the mean VINCULIN or β-ACTIN levels in nuclear fractions normalized to cytoplasmic fractions, and mean HISTONE H3 levels in cytoplasmic fractions normalized to nuclear fractions, from 11 (n=2), 35 (n=2), and 80-week-old (n=2) mice. Data are presented as means ± SD. (b) Volcano plot showing differential subcellular protein distribution between podocytes from mice aged 80 and 35 weeks. The x-axis represents the log2 fold change of the nuclear-to-cytoplasmic (N/C) ratio (80 versus 35 weeks), and the y-axis shows −log10 (P value). Proteins not meeting significance thresholds are shown in gray. (c) Scatter plot showing correlation of subcellular protein distribution between podocytes from mice aged 80 and 35 weeks. The x-axis represents the log2-transformed N/C ratio in 35-week samples, and the y-axis the corresponding values in 80-week samples. Each point represents an individual protein. Solid lines at zero on both axes indicate equal nuclear and cytoplasmic distribution. (d) Volcano plot showing differential subcellular protein distribution between podocytes from mice aged 35 and 11 weeks. The x-axis represents the log2 fold change of the N/C ratio (35 versus 11 weeks), and the y-axis shows −log10 (P value). Proteins not meeting significance thresholds are shown in gray. (e) Scatter plot showing correlation of subcellular protein distribution between podocytes from mice aged 35 and 11 weeks. The x-axis represents the log2-transformed N/C ratio in 11-week samples, and the y-axis the corresponding values in 35-week samples. Each point represents an individual protein. Solid lines at zero on both axes indicate equal nuclear and cytoplasmic distribution.

**Supplemental Figure 2:**
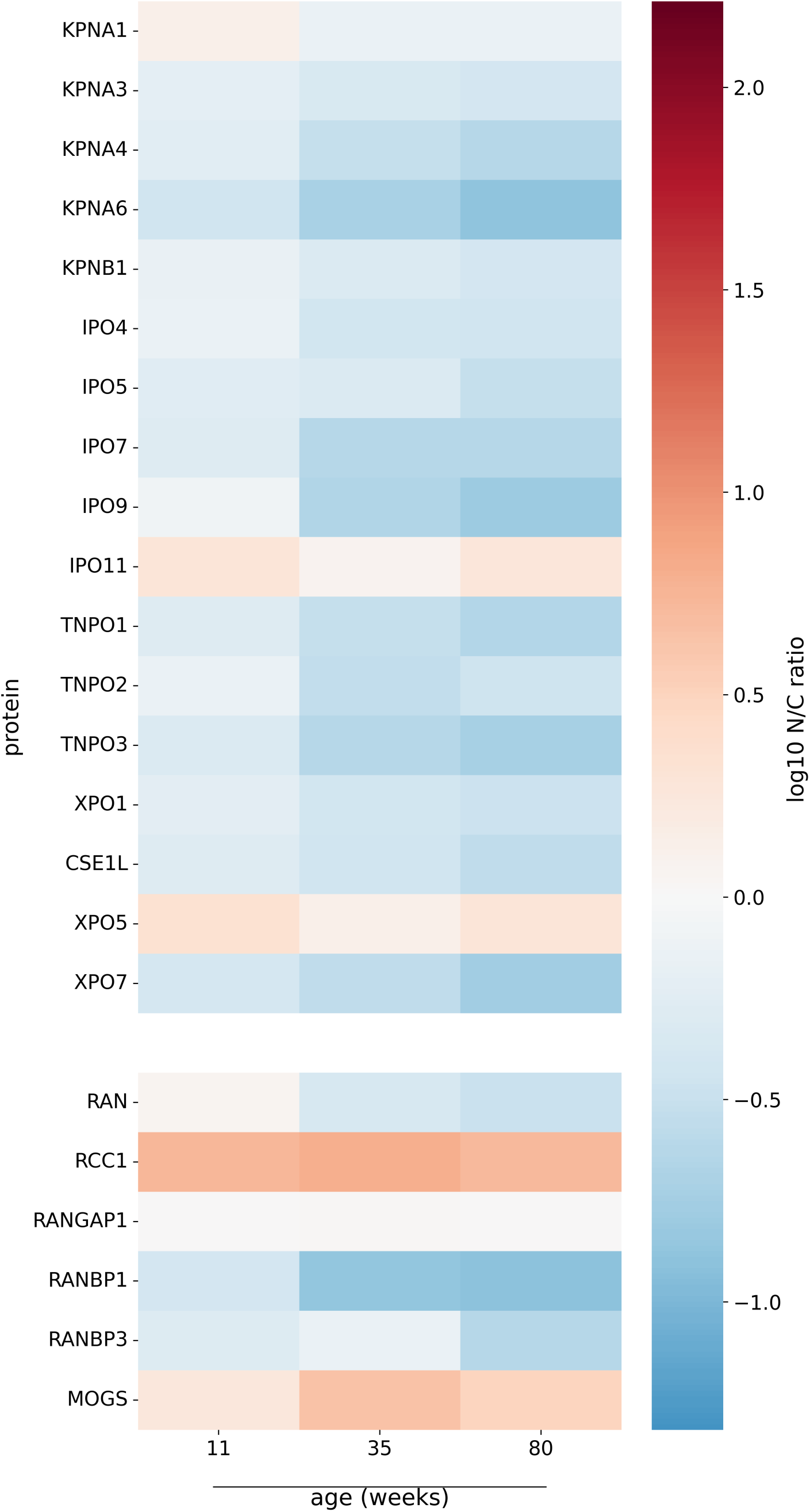
Subcellular distribution of nuclear transport receptors and transport-related factors is largely preserved across aging in podocytes. Heat map showing age-associated changes in subcellular distribution of nuclear transport receptors and transport-related factors in podocytes from mice aged 11, 35, and 80 weeks. Values represent the log2-transformed nuclear-to-cytoplasmic (N/C) ratio for each protein. Data are z-score–normalized across samples; color intensity indicates relative abundance. Each column shows the mean of independent biological replicates (n = 3 for 11 weeks; n = 3 for 35 weeks; n = 2 for 80 weeks).

**Supplemental Figure 3:**
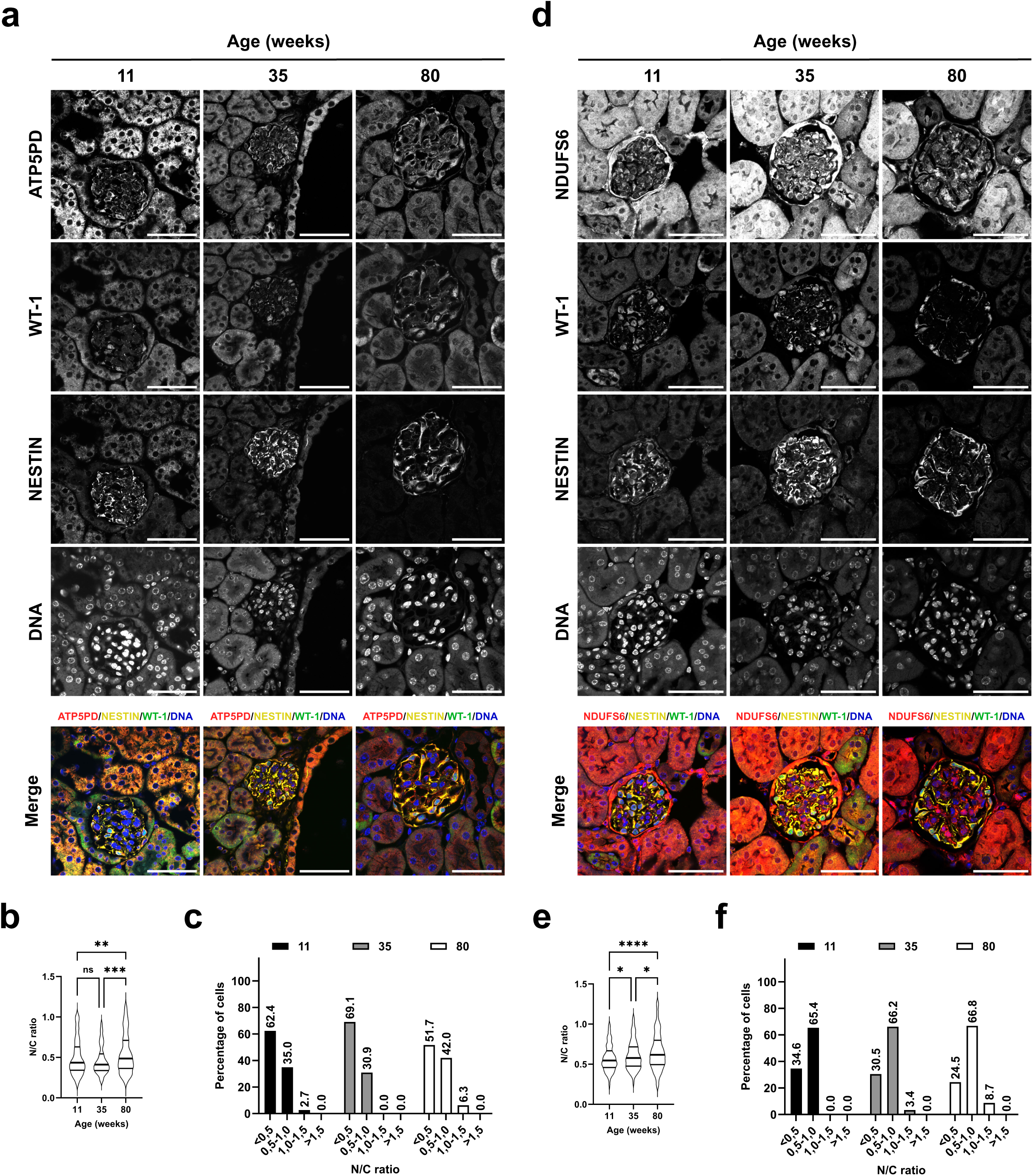
Mitochondrial proteins show cytoplasmic-to-nuclear relocalization during aging. (**a, d**) Confocal images of podocytes stained for ATP5PD (**a**) and NDUFS6 (**d**). NESTIN was used as a podocyte-specific cytoplasmic marker, WT-1 as a podocyte-specific nuclear marker, and DAPI as a nuclear stain. Scale bar, 50 μm. (b) Nuclear-to-cytoplasmic (N/C) fluorescence intensity ratio of ATP5PD, quantified from (**a**) (11 weeks, n = 269 from 6 mice; 35 weeks, n = 165 from 4 mice; 80 weeks, n = 587 from 9 mice). Data are displayed as violin plots with median (thick line) and interquartile range (thin lines). ns, non-significant; **P < 0.01, ***P < 0.001. Kruskal-Wallis non-parametric test, one-way ANOVA, uncorrected Dunn’s test, multiple comparisons; 11 weeks vs. 35 weeks, P=0.1636, 11 weeks vs. 80 weeks, **P=0.0072, 35 weeks vs. 80 weeks, ***P=0.0001. (c) Proportion of podocytes within defined N/C ratio categories (<0.5, 0.5–1.0, 1.0–1.5, >1.5) based on measurements in (**a**), shown for each age group (11, 35, and 80 weeks); percentages are indicated above bars. (e) N/C ratio of NDUFS6 fluorescence intensity quantified from (**d**), based on 244–376 podocytes per condition (11 weeks, n = 244 from 6 mice; 35 weeks, n = 268 from 7 mice; 80 weeks, n = 376 from 8 mice), presented as in (**b**). *P < 0.05, \*\*\**P* < 0.0001. Kruskal-Wallis non-parametric test, one-way ANOVA, uncorrected Dunn’s test, multiple comparisons; 11 weeks vs. 35 weeks, *P=0.0277, 11 weeks vs. 80 weeks, ****P < 0.0001, 35 weeks vs. 80 weeks, *P=0.0240. (f) Distribution of podocytes across N/C ratio categories for NDUFS6, derived from (**d**) and displayed as in (**c**).

**Supplemental Figure 4:**
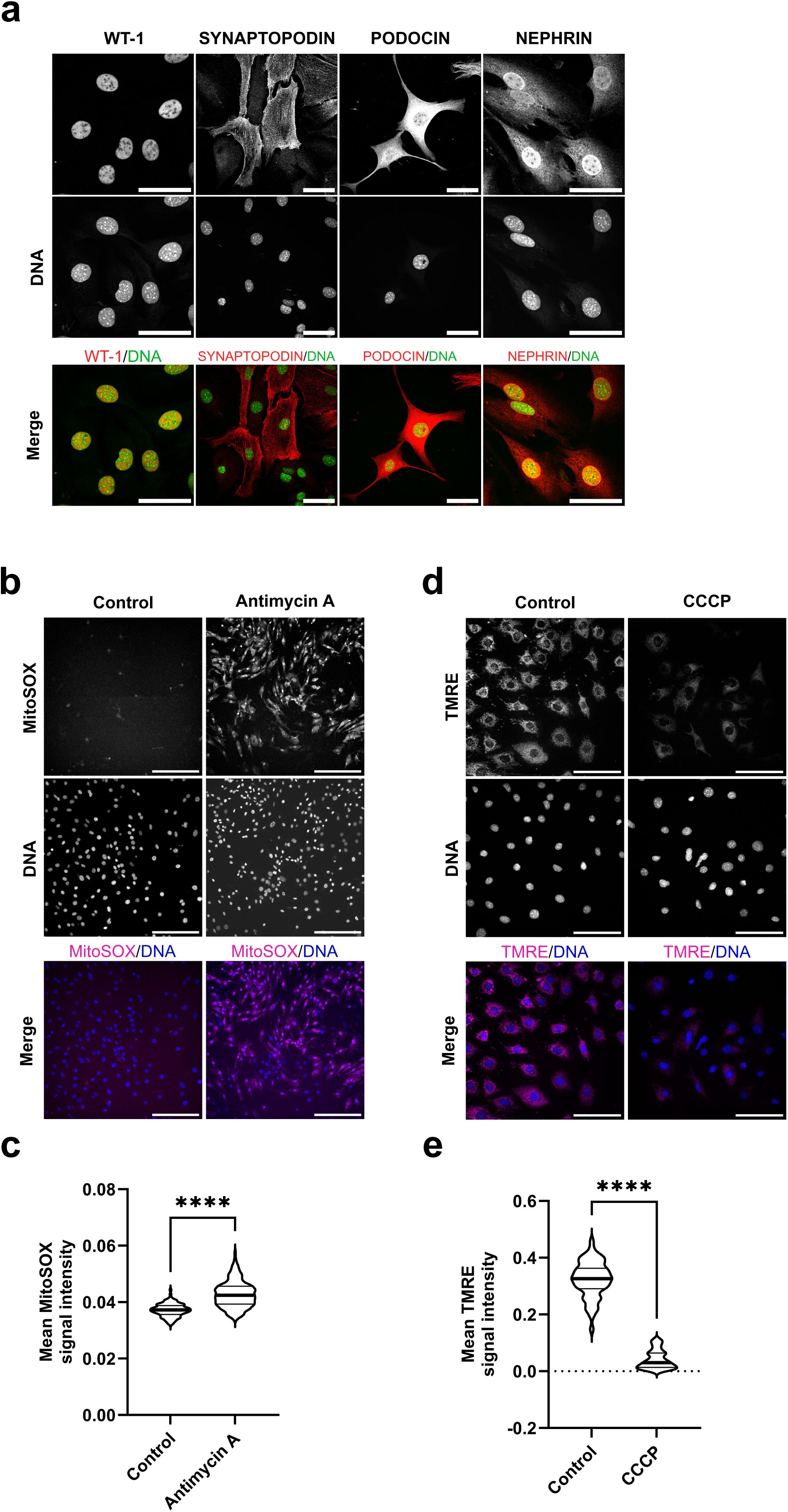
Mitochondrial dysfunction induces ROS production and membrane depolarization in cultured podocytes. (a) Representative immunofluorescence images of cultured podocytes stained with antibodies against WT-1, SYNAPTOPODIN, PODOCIN, and NEPHRIN, DNA visualized with DAPI. Scale bar, 20 µm. (**b, d**) Podocytes were treated with 10 μM antimycin A (**b**) or 25 μM CCCP (**d**) for 6 h, followed by staining with 400 µM MitoSOX or 500 µM TMRE for 30 minutes, respectively. Nuclei were visualized using Hoechst. Scale bar, 200 µm for *B,* 100 µm for *D*. (**c**) Quantification of mean MitoSOX fluorescence intensity based on data from (**b**). n = 419 for control, n = 1371 for antimycin A from three independent experiments. Data are presented as violin plots; thick lines indicate medians and thin lines indicate the 25th and 75th percentiles. \*\*\*\**P*< 0.0001. Unpaired experimental design, nonparametric test, Mann-Whitney test, two-tailed. (**e**) Quantification of mean TMRE fluorescence intensity based on data from (**d**). n = 55 for control, n = 54 for CCCP from three independent experiments. Thick lines indicate medians, and thin lines indicate the 25th and 75th percentiles. \*\*\*\**P*< 0.0001. Unpaired experimental design, nonparametric test, Mann-Whitney test, two-tailed.

**Supplemental Figure 5:**
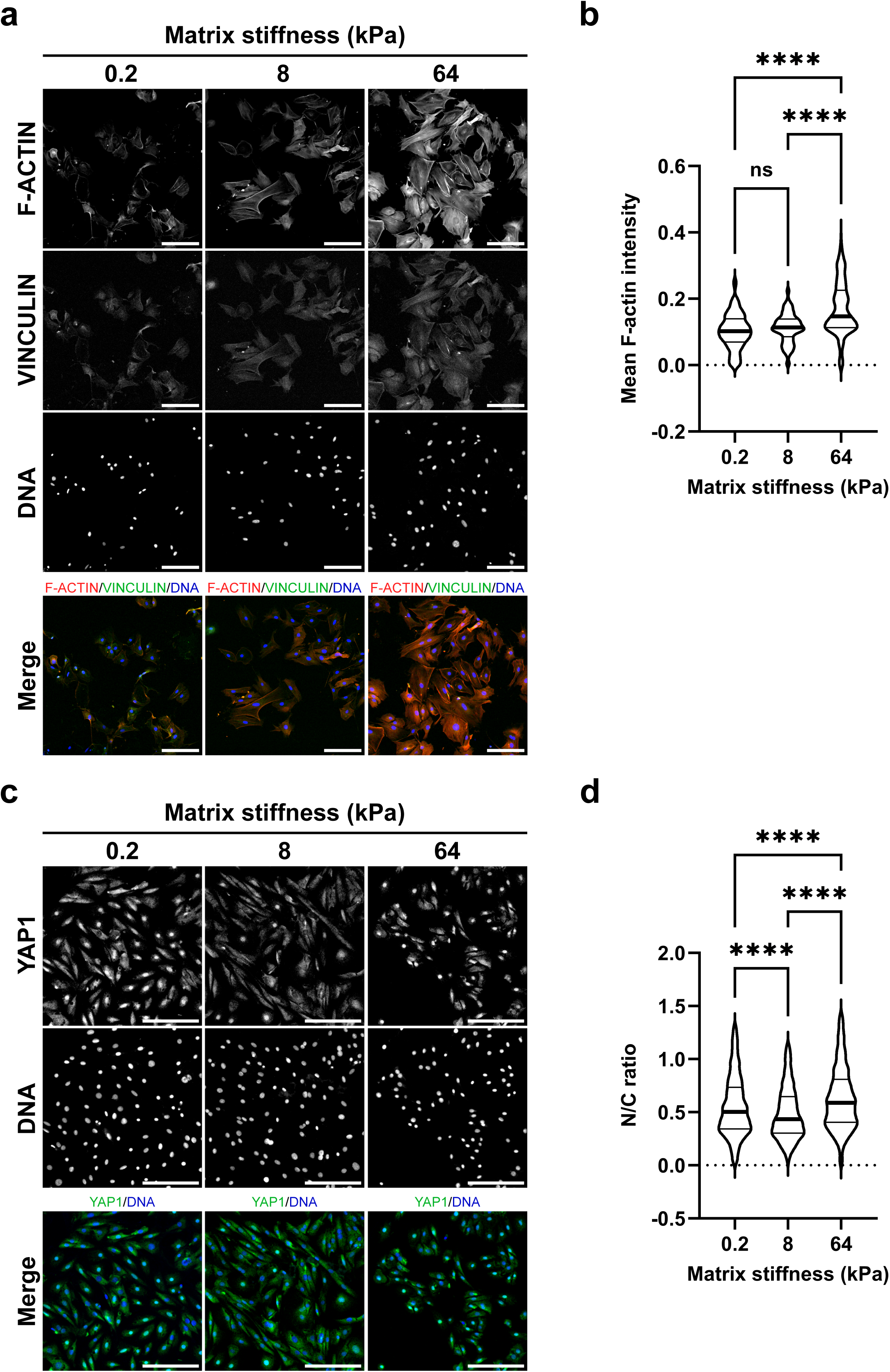
Matrix stiffness modulates stress fiber formation and YAP1 localization in podocytes. (**a, c**) Representative images of podocytes cultured on substrates of defined stiffness (0.2, 8, and 64 kPa). Cells were stained for F-ACTIN and VINCULIN (**a**) or YAP1 (**c**), with nuclei visualized using DAPI. Scale bar, 200 μm. (b) Quantification of mean F-ACTIN fluorescence intensity based on data from (**a**) (n = 191, 0.2 kPa; n = 242, 8 kPa; n = 276, 64 kPa cells from three independent experiments). Median (thick line) and interquartile range (thin lines). ns, non-significant, ****P< 0.0001. Kruskal-Wallis non-parametric test, one-way ANOVA, uncorrected Dunn’s test, multiple comparisons; 0.2 kPa vs. 8 kPa, P < 0.0001, 0.2 kPa vs. 64 kPa, P < 0.0001, 8 kPa vs. 64 kPa, P=0.0979. (**d**) Quantification of the nuclear-to-cytoplasmic (N/C) ratio of YAP1 based on data from (**c**) (n = 1475, 0.2 kPa; n = 1224, 8 kPa; n = 778, 64 kPa cells from three independent experiments). ****P< 0.0001. Kruskal-Wallis non-parametric test, one-way ANOVA, uncorrected Dunn’s test, multiple comparisons.

**Supplemental Figure 6:**
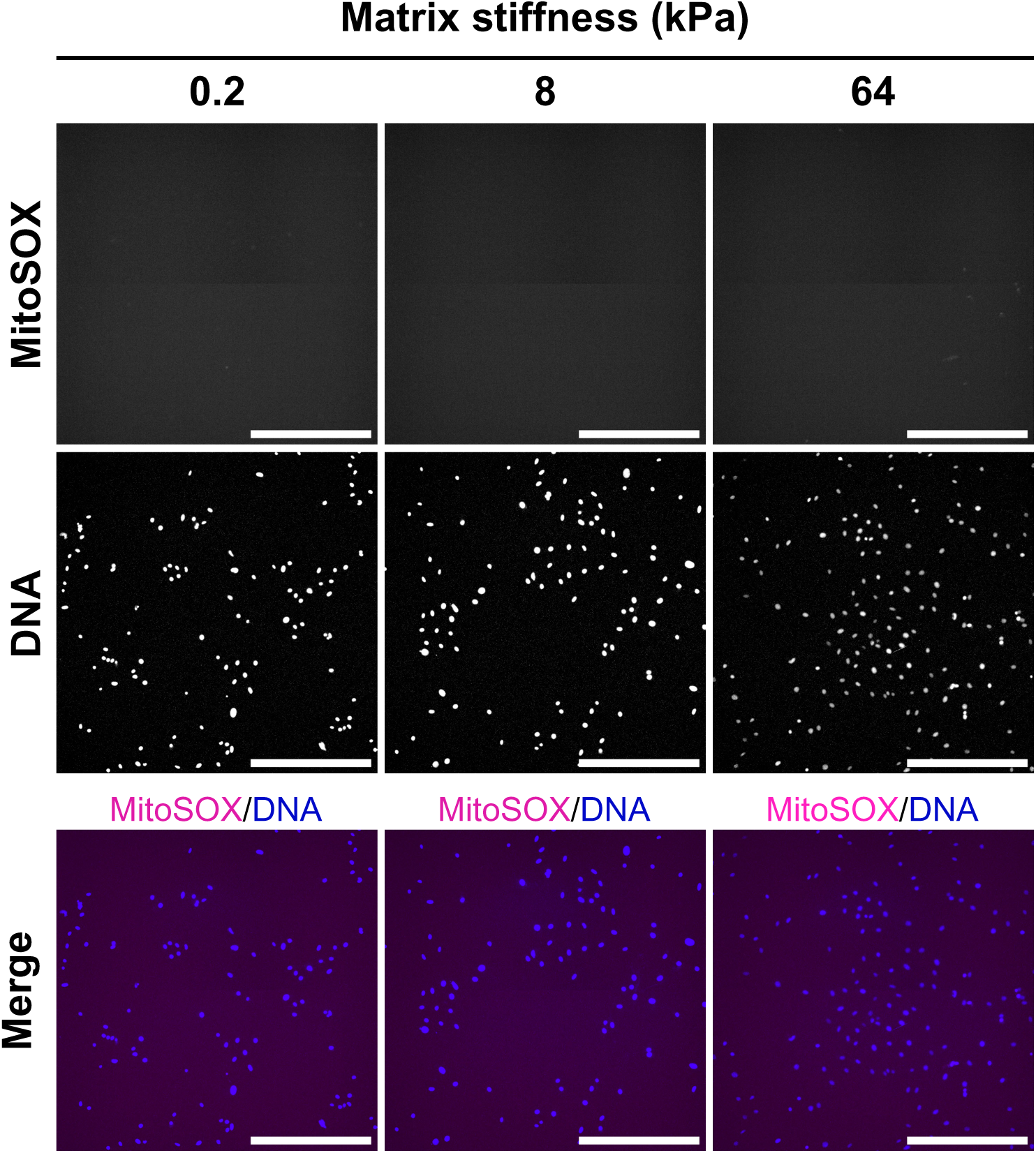
Matrix stiffness does not induce superoxide production. Representative images of podocytes cultured on substrates of defined stiffness (0.2, 8, and 64 kPa) and stained with MitoSOX to assess superoxide production. Nuclei were visualized using Hoechst 33342. Scale bar, 200 μm.

**Supplemental table 1:**
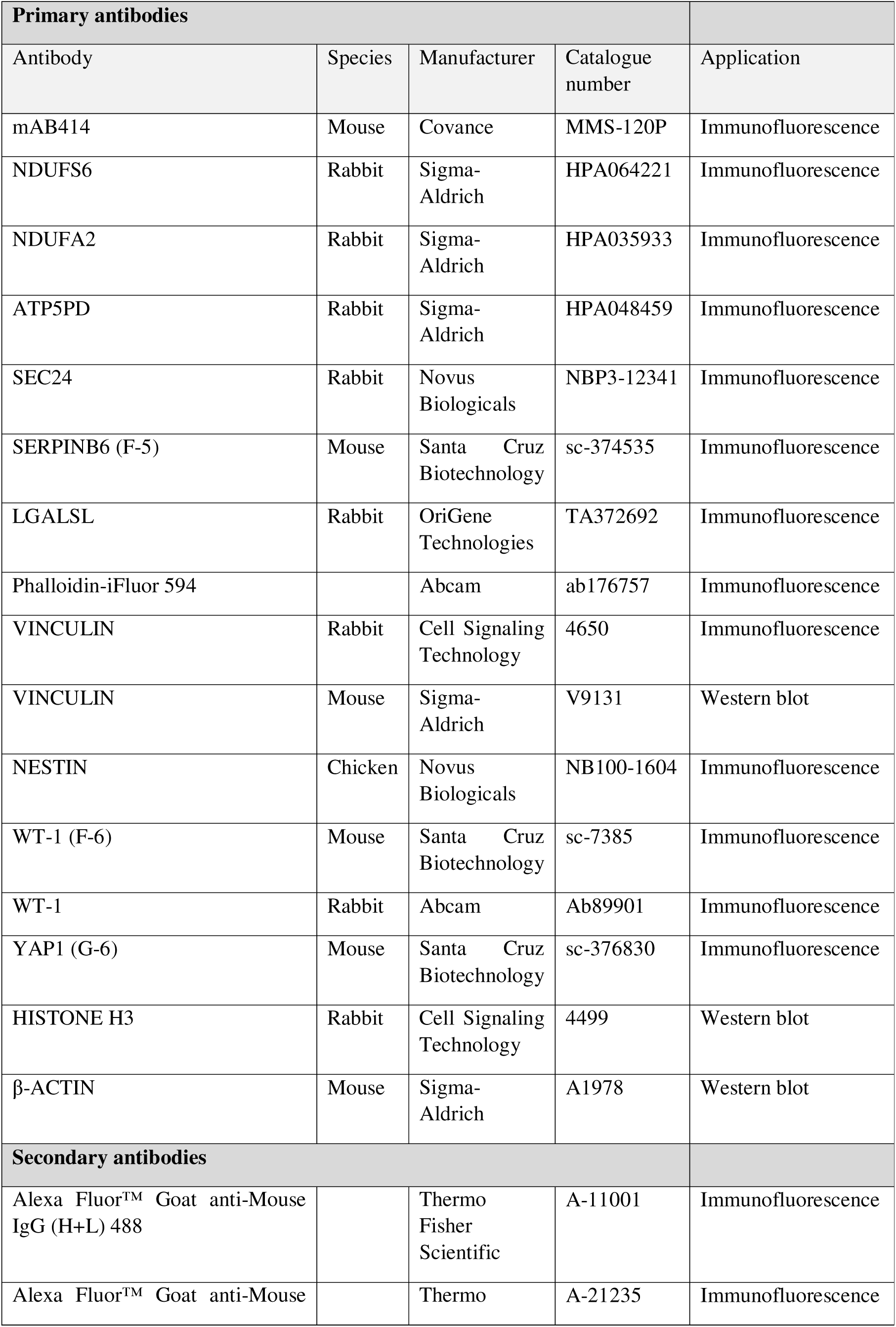
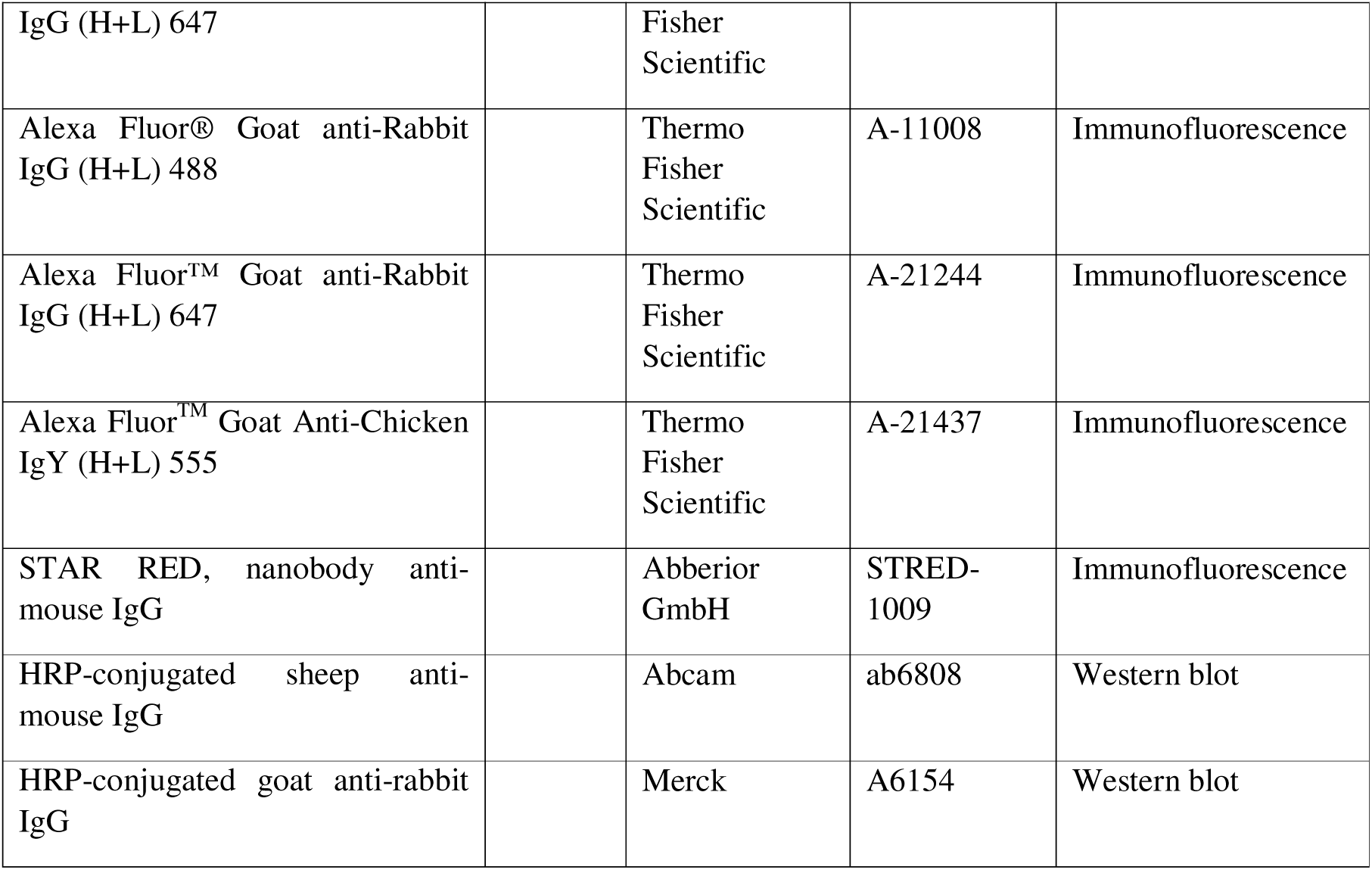
List of antibodies.

## References

1. de Duve C. The origin of eukaryotes: a reappraisal. Nat Rev Genet. 2007;8(5):395–403.

2. Bar-Peled L, Kory N. Principles and functions of metabolic compartmentalization. Nat Metab. 2022;4(10):1232–44.

3. Pawson T, Scott JD. Signaling through scaffold, anchoring, and adaptor proteins. Science. 1997;278(5346):2075–80.

4. De Magistris P, Antonin W. The Dynamic Nature of the Nuclear Envelope. Curr Biol. 2018;28(8):R487–R97.

5. Sigaeva A, Hutchings C, Cesnik A, Lilley KS, Lundberg E. Subcellular localization as a driver of protein function. Nat Rev Mol Cell Biol. 2026.

6. Wu M, Tao H, Xu T, Zheng X, Wen C, Wang G, et al. Spatial proteomics: unveiling the multidimensional landscape of protein localization in human diseases. Proteome Sci. 2024;22(1):7.

7. Hung MC, Link W. Protein localization in disease and therapy. J Cell Sci. 2011;124(Pt 20):3381–92.

8. Yankner BA, Lu T, Loerch P. The aging brain. Annu Rev Pathol. 2008;3:41–66.

9. Shankland SJ, Rule AD, Kutz JN, Pippin JW, Wessely O. Podocyte Senescence and Aging. Kidney360. 2023;4(12):1784–93.

10. Nagata M. Podocyte injury and its consequences. Kidney Int. 2016;89(6):1221–30.

11. Shankland SJ, Wang Y, Shaw AS, Vaughan JC, Pippin JW, Wessely O. Podocyte Aging: Why and How Getting Old Matters. J Am Soc Nephrol. 2021;32(11):2697–713.

12. Puelles VG, Cullen-McEwen LA, Taylor GE, Li J, Hughson MD, Kerr PG, et al. Human podocyte depletion in association with older age and hypertension. Am J Physiol Renal Physiol. 2016;310(7):F656–F68.

13. Wharram BL, Goyal M, Wiggins JE, Sanden SK, Hussain S, Filipiak WE, et al. Podocyte depletion causes glomerulosclerosis: diphtheria toxin-induced podocyte depletion in rats expressing human diphtheria toxin receptor transgene. Journal of the American Society of Nephrology : JASN. 2005;16(10):2941–52.

14. Wente SR, Rout MP. The Nuclear Pore Complex and Nuclear Transport. Cold Spring Harbor Perspectives in Biology. 2010;2(10):a000562-a.

15. Gorlich D, Kutay U. Transport between the cell nucleus and the cytoplasm. Annu Rev Cell Dev Biol. 1999;15:607–60.

16. Timney BL, Raveh B, Mironska R, Trivedi JM, Kim SJ, Russel D, et al. Simple rules for passive diffusion through the nuclear pore complex. J Cell Biol. 2016;215(1):57–76.

17. Dutta S, Sengupta P. Men and mice: Relating their ages. Life Sci. 2016;152:244–8.

18. Kikuchi M, Wickman L, Rabah R, Wiggins RC. Podocyte number and density changes during early human life. Pediatric nephrology (Berlin, Germany). 2017;32(5):823–34.

19. D’Angelo MA, Raices M, Panowski SH, Hetzer MW. Age-dependent deterioration of nuclear pore complexes causes a loss of nuclear integrity in postmitotic cells. Cell. 2009;136(2):284–95.

20. Braun DA, Sadowski CE, Kohl S, Lovric S, Astrinidis SA, Pabst WL, et al. Mutations in nuclear pore genes NUP93, NUP205 and XPO5 cause steroid-resistant nephrotic syndrome. Nature Genetics. 2016;48(4):457–65.

21. Braun DA, Lovric S, Schapiro D, Schneider R, Marquez J, Asif M, et al. Mutations in multiple components of the nuclear pore complex cause nephrotic syndrome. Journal of Clinical Investigation. 2018;128(10):4313–28.

22. Davis LI, Blobel G. Identification and characterization of a nuclear pore complex protein. Cell. 1986;45(5):699–709.

23. Agborbesong E, Zhou JX, Li LX, Calvet JP, Li X. Antioxidant enzyme peroxiredoxin 5 regulates cyst growth and ciliogenesis via modulating Plk1 stability. FASEB J. 2022;36(1):e22089.

24. Elshani M, Um IH, Leung S, Reynolds PA, Chapman A, Kudsy M, et al. Transcription Factor NFE2L1 Decreases in Glomerulonephropathies after Podocyte Damage. Cells. 2023;12(17).

25. Boye TL, Jeppesen JC, Maeda K, Pezeshkian W, Solovyeva V, Nylandsted J, et al. Annexins induce curvature on free-edge membranes displaying distinct morphologies. Sci Rep. 2018;8(1):10309.

26. Song L, Shen W, Wang L, Song J, Tu W, Ke B, et al. Annexin A1 may contribute to the morphological changes in podocytes by mediating endocytic vesicle fusion and transport via promotion of SNARE assembly in idiopathic membranous nephropathy. Nephrology (Carlton). 2024;29(2):76–85.

27. Yasuda H, Fukusumi Y, Zhang Y, Kawachi H. 14-3-3 Proteins stabilize actin and vimentin filaments to maintain processes in renal glomerular podocyte. FASEB J. 2023;37(10):e23168.

28. Garg P, Verma R, Cook L, Soofi A, Venkatareddy M, George B, et al. Actin-depolymerizing factor cofilin-1 is necessary in maintaining mature podocyte architecture. J Biol Chem. 2010;285(29):22676–88.

29. Gao Y, Hamed M, Martin IV, Raffetseder U, Liu X, Leitz A, et al. The nuclear export inhibitor selinexor improves kidney function in a rat model of focal segmental glomerulosclerosis. Am J Physiol Renal Physiol. 2025;329(4):F496–f509.

30. Nakata H, Yamamoto M, Kumchantuek T, Adhapanyawanich K, Nishiuchi T, Iseki S. Synthesis, localization and possible function of serine (or cysteine) peptidase inhibitor, clade B, member 6a (Serpinb6a) in mouse submandibular gland. Cell Tissue Res. 2017;369(3):513–26.

31. Bhat R, Belardi B, Mori H, Kuo P, Tam A, Hines WC, et al. Nuclear repartitioning of galectin-1 by an extracellular glycan switch regulates mammary morphogenesis. Proc Natl Acad Sci U S A. 2016;113(33):E4820–7.

32. Adams EJ, Chen XW, O’Shea KS, Ginsburg D. Mammalian COPII coat component SEC24C is required for embryonic development in mice. J Biol Chem. 2014;289(30):20858–70.

33. Liu Q, Chang CE, Wooldredge AC, Fong B, Kennedy BK, Zhou C. Tom70-based transcriptional regulation of mitochondrial biogenesis and aging. Elife. 2022;11.

34. Khan AH, Gu X, Patel RJ, Chuphal P, Viana MP, Brown AI, et al. Mitochondrial protein heterogeneity stems from the stochastic nature of co-translational protein targeting in cell senescence. Nat Commun. 2024;15(1):8274.

35. Wang Y, Eng DG, Kaverina NV, Loretz CJ, Koirala A, Akilesh S, et al. Global transcriptomic changes occur in aged mouse podocytes. Kidney Int. 2020;98(5):1160–73.

36. Rosa FLL, de Souza IIA, Monnerat G, Campos de Carvalho AC, Maciel L. Aging Triggers Mitochondrial Dysfunction in Mice. Int J Mol Sci. 2023;24(13).

37. Espinoza K, Schaler AW, Gray DT, Sass AR, Escobar A, Moore K, et al. Dynamic changes in mitochondria support phenotypic flexibility of microglia. Nat Commun. 2025;16(1):11103.

38. Esposito LA, Melov S, Panov A, Cottrell BA, Wallace DC. Mitochondrial disease in mouse results in increased oxidative stress. Proc Natl Acad Sci U S A. 1999;96(9):4820–5.

39. Witte S, Boshnakovska A, Özdemir M, Chowdhury A, Rehling P, Aich A. Defective COX1 expression in aging mice liver. Biol Open. 2023;12(3).

40. Kabgani N, Grigoleit T, Schulte K, Sechi A, Sauer-Lehnen S, Tag C, et al. Primary cultures of glomerular parietal epithelial cells or podocytes with proven origin. PLoS One. 2012;7(4):e34907.

41. Smeets B, Kabgani N, Moeller MJ. Isolation and Primary Culture of Murine Podocytes with Proven Origin. Methods Mol Biol. 2016;1397:3–10.

42. Quinlan CL, Gerencser AA, Treberg JR, Brand MD. The mechanism of superoxide production by the antimycin-inhibited mitochondrial Q-cycle. J Biol Chem. 2011;286(36):31361–72.

43. Zhang YQ, Shen X, Xiao XL, Liu MY, Li SL, Yan J, et al. Mitochondrial uncoupler carbonyl cyanide m-chlorophenylhydrazone induces vasorelaxation without involving K. Br J Pharmacol. 2016;173(21):3145–58.

44. Liao XD, Wang XH, Jin HJ, Chen LY, Chen Q. Mechanical stretch induces mitochondria-dependent apoptosis in neonatal rat cardiomyocytes and G2/M accumulation in cardiac fibroblasts. Cell Res. 2004;14(1):16–26.

45. Ke W, Wang B, Liao Z, Song Y, Li G, Ma L, et al. Matrix stiffness induces Drp1-mediated mitochondrial fission through Piezo1 mechanotransduction in human intervertebral disc degeneration. J Transl Med. 2023;21(1):711.

46. Zhang Z, Cui S, Fu Y, Wang J, Liu J, Wei F. Mechanical force induces mitophagy-mediated anaerobic oxidation in periodontal ligament stem cells. Cell Mol Biol Lett. 2023;28(1):57.

47. Lennon R, Randles MJ, Humphries MJ. The importance of podocyte adhesion for a healthy glomerulus. Front Endocrinol (Lausanne). 2014;5:160.

48. Haydak J, Azeloglu EU. Role of biophysics and mechanobiology in podocyte physiology. Nat Rev Nephrol. 2024;20(6):371–85.

49. Hu M, Azeloglu EU, Ron A, Tran-Ba KH, Calizo RC, Tavassoly I, et al. A biomimetic gelatin-based platform elicits a pro-differentiation effect on podocytes through mechanotransduction. Sci Rep. 2017;7:43934.

50. Zhang Y, Musah S. Mechanosensitive Differentiation of Human iPS Cell-Derived Podocytes. Bioengineering (Basel). 2024;11(10).

51. Ma SJ, Zhu YT, He FF, Zhang C. Mechanisms and Therapeutic Perspectives of Podocyte Aging in Podocytopathies. Int J Mol Sci. 2025;26(18).

52. Zhao Q, Huang Y, Fu N, Cui C, Peng X, Kang H, et al. Podocyte senescence: from molecular mechanisms to therapeutics. Ren Fail. 2024;46(2):2398712.

53. Kutay U, Juhlen R, Antonin W. Mitotic disassembly and reassembly of nuclear pore complexes. Trends Cell Biol. 2021;31(12):1019–33.

54. Rempel IL, Crane MM, Thaller DJ, Mishra A, Jansen DP, Janssens G, et al. Age-dependent deterioration of nuclear pore assembly in mitotic cells decreases transport dynamics. eLife. 2019;8.

55. Toyama Brandon H, Savas Jeffrey N, Park Sung K, Harris Michael S, Ingolia Nicholas T, Yates John R, et al. Identification of Long-Lived Proteins Reveals Exceptional Stability of Essential Cellular Structures. Cell. 2013;154(5):971–82.

56. Kim HJ, Taylor JP. Lost in Transportation: Nucleocytoplasmic Transport Defects in ALS and Other Neurodegenerative Diseases. Neuron. 2017;96(2):285–97.

57. Hutten S, Dormann D. Nucleocytoplasmic transport defects in neurodegeneration - Cause or consequence? Semin Cell Dev Biol. 2020;99:151–62.

58. Khalil B, Linsenmeier M, Smith CL, Shorter J, Rossoll W. Nuclear-import receptors as gatekeepers of pathological phase transitions in ALS/FTD. Mol Neurodegener. 2024;19(1):8.

59. Chen PX, Zhang L, Wu X, Liang Z, Hao X, Zhu Q, et al. Mitochondrial superoxide regulates nuclear envelope integrity and ageing via redox-mediated lipid metabolism. Nat Metab. 2026;8(2):371–88.

60. Chen G, Dong H, Tian Y. Mito-nuclear communication: From cellular responses to organismal health. Mol Cell. 2026;86(3):522–32.

61. Szczesny B, Hazra TK, Papaconstantinou J, Mitra S, Boldogh I. Age-dependent deficiency in import of mitochondrial DNA glycosylases required for repair of oxidatively damaged bases. Proc Natl Acad Sci U S A. 2003;100(19):10670–5.

62. Bogorodskiy A, Okhrimenko I, Burkatovskii D, Jakobs P, Maslov I, Gordeliy V, et al. Role of Mitochondrial Protein Import in Age-Related Neurodegenerative and Cardiovascular Diseases. Cells. 2021;10(12).

63. Nargund AM, Pellegrino MW, Fiorese CJ, Baker BM, Haynes CM. Mitochondrial import efficiency of ATFS-1 regulates mitochondrial UPR activation. Science. 2012;337(6094):587–90.

64. Bratic A, Larsson NG. The role of mitochondria in aging. J Clin Invest. 2013;123(3):951–7.

65. Sun N, Youle RJ, Finkel T. The Mitochondrial Basis of Aging. Mol Cell. 2016;61(5):654–66.

66. Yousef A, Fang L, Heidari M, Huang A, Kondraciuk P, Yee KA, et al. Alterations in mitochondria and cellular senescence in aged sEH null female kidneys. Geroscience. 2026;48(2):2707–25.

67. Galvan DL, Green NH, Danesh FR. The hallmarks of mitochondrial dysfunction in chronic kidney disease. Kidney Int. 2017;92(5):1051–7.

68. Duann P, Lin PH. Mitochondria Damage and Kidney Disease. Adv Exp Med Biol. 2017;982:529–51.

69. Brinkkoetter PT, Bork T, Salou S, Liang W, Mizi A, Özel C, et al. Anaerobic Glycolysis Maintains the Glomerular Filtration Barrier Independent of Mitochondrial Metabolism and Dynamics. Cell Rep. 2019;27(5):1551–66.e5.

70. Daehn I, Casalena G, Zhang T, Shi S, Fenninger F, Barasch N, et al. Endothelial mitochondrial oxidative stress determines podocyte depletion in segmental glomerulosclerosis. J Clin Invest. 2014;124(4):1608–21.

71. Özel C, Reitmeier KM, Kieckhöfer E, Abualia K, Nguyen-Minh D, Matin M, et al. Disruption of Mitochondrial Dynamics and Integrity Drives Divergent Metabolic Flexibility and Resilience in Podocytes. FASEB J. 2025;39(24):e71340.

72. Aragona M, Panciera T, Manfrin A, Giulitti S, Michielin F, Elvassore N, et al. A mechanical checkpoint controls multicellular growth through YAP/TAZ regulation by actin-processing factors. Cell. 2013;154(5):1047–59.

73. Dupont S, Morsut L, Aragona M, Enzo E, Giulitti S, Cordenonsi M, et al. Role of YAP/TAZ in mechanotransduction. Nature. 2011;474(7350):179–83.

74. Sorrentino G, Ruggeri N, Specchia V, Cordenonsi M, Mano M, Dupont S, et al. Metabolic control of YAP and TAZ by the mevalonate pathway. Nat Cell Biol. 2014;16(4):357–66.

75. DeRan M, Yang J, Shen CH, Peters EC, Fitamant J, Chan P, et al. Energy stress regulates hippo-YAP signaling involving AMPK-mediated regulation of angiomotin-like 1 protein. Cell Rep. 2014;9(2):495–503.

76. Mo JS, Meng Z, Kim YC, Park HW, Hansen CG, Kim S, et al. Cellular energy stress induces AMPK-mediated regulation of YAP and the Hippo pathway. Nat Cell Biol. 2015;17(4):500–10.

77. Szeto SG, Narimatsu M, Lu M, He X, Sidiqi AM, Tolosa MF, et al. YAP/TAZ Are Mechanoregulators of TGF-. J Am Soc Nephrol. 2016;27(10):3117–28.

78. Chen J, Wang X, He Q, Bulus N, Fogo AB, Zhang MZ, et al. YAP Activation in Renal Proximal Tubule Cells Drives Diabetic Renal Interstitial Fibrogenesis. Diabetes. 2020;69(11):2446–57.

79. Xu J, Li PX, Wu J, Gao YJ, Yin MX, Lin Y, et al. Involvement of the Hippo pathway in regeneration and fibrogenesis after ischaemic acute kidney injury: YAP is the key effector. Clin Sci (Lond). 2016;130(5):349–63.

80. Endlich N, Kress KR, Reiser J, Uttenweiler D, Kriz W, Mundel P, et al. Podocytes respond to mechanical stress in vitro. J Am Soc Nephrol. 2001;12(3):413–22.

81. Charras G, Sahai E. Physical influences of the extracellular environment on cell migration. Nat Rev Mol Cell Biol. 2014;15(12):813–24.

82. Abrass CK, Adcox MJ, Raugi GJ. Aging-associated changes in renal extracellular matrix. Am J Pathol. 1995;146(3):742–52.

83. Wiggins JE. Aging in the glomerulus. J Gerontol A Biol Sci Med Sci. 2012;67(12):1358–64.

84. Yao L, Li Y, Wang P, Xu C, Yu Z. Nucleoporin-associated steroid-resistant nephrotic syndrome. Pediatr Nephrol. 2025;40(3):629–49.

85. Lee JH, Tingey M, Zhang Z, Buerger F, Hong J, Zhang G, et al. N. Nat Cell Biol. 2026;28(3):553–66.

86. Puelles VG, Fleck D, Ortz L, Papadouri S, Strieder T, Böhner AMC, et al. Novel 3D analysis using optical tissue clearing documents the evolution of murine rapidly progressive glomerulonephritis. Kidney Int. 2019;96(2):505–16.

87. Hughes CS, Foehr S, Garfield DA, Furlong EE, Steinmetz LM, Krijgsveld J. Ultrasensitive proteome analysis using paramagnetic bead technology. Mol Syst Biol. 2014;10(10):757.

88. Thompson A, Wölmer N, Koncarevic S, Selzer S, Böhm G, Legner H, et al. TMTpro: Design, Synthesis, and Initial Evaluation of a Proline-Based Isobaric 16-Plex Tandem Mass Tag Reagent Set. Anal Chem. 2019;91(24):15941–50.

89. Yang F, Shen Y, Camp DG, Smith RD. High-pH reversed-phase chromatography with fraction concatenation for 2D proteomic analysis. Expert Rev Proteomics. 2012;9(2):129–34.

90. Kong AT, Leprevost FV, Avtonomov DM, Mellacheruvu D, Nesvizhskii AI. MSFragger: ultrafast and comprehensive peptide identification in mass spectrometry-based proteomics. Nat Methods. 2017;14(5):513–20.

91. Huber W, von Heydebreck A, Sültmann H, Poustka A, Vingron M. Variance stabilization applied to microarray data calibration and to the quantification of differential expression. Bioinformatics. 2002;18 Suppl 1:S96–104.

92. Ritchie ME, Phipson B, Wu D, Hu Y, Law CW, Shi W, et al. limma powers differential expression analyses for RNA-sequencing and microarray studies. Nucleic Acids Res. 2015;43(7):e47.

